# SLX4IP acts in parallel to FANCM to limit BLM-dependent replication stress at ALT telomeres

**DOI:** 10.1101/2025.05.28.656696

**Authors:** Jessica Spindler, Francesca Pandolfo, Anna Eva Koch, Priscilla Piccirillo, Johanna Bihler, Marcel Morgenstern, Sandra Buschbaum, Joshua Coon, Robert Hänsel-Hertsch, Kavi P. M. Mehta, Stephanie Panier

## Abstract

Alternative Lengthening of Telomeres (ALT) is a telomerase-independent telomere maintenance mechanism that enables cancer cells to gain unlimited replicative capacity. ALT relies on recombination-mediated telomere elongation and is promoted by telomeric replication stress. However, ALT requires strict regulation, as excessive replication stress or recombination are cytotoxic. Central to ALT is the RecQ helicase BLM, which regulates telomeric replication stress and promotes telomere recombination and DNA synthesis. Despite its key role in the ALT pathway, BLM must be tightly regulated to prevent deleterious outcomes. Here, we identify SLX4IP as a key suppressor of BLM-driven replication stress at ALT telomeres. Loss of SLX4IP in ALT-positive cells leads to BLM-dependent telomeric replication stress and impaired replication fork progression. Mechanistically, SLX4IP limits the unwinding of unligated Okazaki fragments by BLM on the lagging strand during telomere replication. This reduces the formation of toxic 5′ DNA flaps and prevents hyperactivation of ATR signalling and deleterious recombination levels. We also uncover a synthetic lethal interaction between SLX4IP and FANCM, an ATPase/translocase that is a known regulator of BLM at telomeric replication forks in ALT cells. We demonstrate that SLX4IP and FANCM act in parallel to restrain BLM activity, thereby maintaining the balance of replication stress and recombination that is necessary for productive ALT. These findings reveal a vulnerability in ALT-positive cancers lacking SLX4IP and establish SLX4IP as a potential biomarker for therapeutic strategies targeting FANCM.

**HIGHLIGHTS:**

- SLX4IP depletion activates the replication stress response at ALT telomeres
- SLX4IP acts in parallel to FANCM to limit replication stress at ALT telomeres
- The synthetic lethal interaction between SLX4IP and FANCM is dependent on BLM
- SLX4IP depletion causes BLM-dependent lagging-strand replication stress

## INTRODUCTION

Telomere maintenance is critical for achieving unlimited replicative potential, which is a defining hallmark of all cancers^1^. While most cancers rely on telomerase to maintain telomere length, approximately 5-15% of cancers, particularly those of mesenchymal origin, employ Alternative Lengthening of Telomeres (ALT) instead^2–4^. Unlike telomerase, which extends telomeres via reverse transcription, ALT utilizes recombination-based pathways to elongate telomeres, with break-induced replication (BIR) playing a central role in this process^5,6^. Consequently, ALT-positive cancer cells co-opt the cellular replication, DNA damage response and recombination machineries to initiate and regulate telomere recombination^6^.

The exact mechanisms underlying ALT activation remain unclear, but it is thought to arise as an adaptive response to the inability to activate telomerase during cellular transformation^7–9^. A key feature of ALT-positive cancer cells is the presence of sustained replication stress at telomeres^10–13^. This replication stress is driven by genetic and epigenetic changes in the telomeric chromatin environment, which render telomeres more permissive to forming G-quadruplexes, TERRA-mediated R-loops, single-strand breaks and oxidative lesions such as 8-oxoguanine. These obstacles impede replication fork progression^14,15^, leading to the activation of ATR-dependent stress signalling pathways^16–19^. A particularly vulnerable aspect of ALT telomere replication is lagging-strand DNA synthesis, where defects in Okazaki fragment maturation result in toxic 5’ DNA flaps and 5’ single-stranded DNA gaps that trigger ATR-dependent replication stress signaling and chromatin PARylation^20,21^.

Telomeric replication stress is essential for generating recombination-prone DNA intermediates that drive template-directed DNA synthesis in ALT-positive cells^15^. However, replication and recombination levels must also be tightly controlled to prevent the accumulation of excessive DNA damage and cytotoxic recombination intermediates, including catenated telomeric DNA^18,19,22–25^. Thus, ALT-positive cells must carefully balance replication stress and recombination levels to ensure productive telomere extension while avoiding cytotoxic levels of genome instability^6^.

A key facilitator of ALT is the RecQ helicase BLM, which plays multiple roles in ALT induction and maintenance^7^. While BLM localizes to both ALT and non-ALT telomeres, it is particularly enriched at ALT telomeres^26,27^. Here, it contributes to ALT-associated PML body (APB) formation through SUMO-mediated liquid-liquid phase separation, creating a microenvironment that facilitates recombination reactions between telomeres^28–30^. BLM also assists in the assembly of the PCNA-RFC-Pol8 replisome at ALT telomeres and promotes branch migration during telomeric DNA synthesis^20,25,31,32^. Additionally, it resolves G-quadruplex structures that accumulate on the G-rich lagging-strand during telomere replication^33,34^ and processes recombination intermediates as part of the BTR dissolvasome complex^35–38^. The BTR complex also helps to rescue stalled replication forks through replication fork regression followed by fork restart^24,39,40^.

Given its multiple roles in ALT, productive ALT telomere extension critically depends on BLM. Indeed, BLM deficiency impairs APB formation and causes telomere shortening^11,28^. However, unrestrained BLM activity results in hyper-ALT phenotypes characterized by elevated replicative stress, particularly on the lagging-strand, excessive DNA synthesis and unprocessed recombination intermediates, all of which compromise cellular viability^20,23–25,41^. Therefore, precise regulation of BLM is critical for ALT-positive cells, as both insufficient and excessive activity can jeopardize telomere maintenance and cell survival.

ALT-positive cells exert significant effort to restrict BLM activity, reducing replication stress and recombination rates to levels that support productive telomere synthesis without impairing cellular viability^42^. One key regulator of BLM is FANCM, an ATPase translocase that limits BLM-dependent excessive replication stress by resolving persistent R-loops and facilitating replication fork remodelling^18,19,22,24^. Importantly, FANCM carries out this function independently of its role within the Fanconi Anemia core complex by directly binding to BLM^24^. Disruption of this interaction is selectively lethal to ALT-positive cells^24^. In parallel, the nuclease DNA2 limits BLM-driven lagging-strand replication stress by removing 5’flaps that are generated by aberrant, BLM-mediated unwinding of unligated Okazaki fragments^20^. In addition, MutSα counteracts BLM activity, likely by promoting heteroduplex rejection and thus preventing premature BLM-dependent branch migration during ALT telomere synthesis^43^. Finally, at the level of recombination intermediates, BLM-mediated dissolution is counteracted by the SLX4-containing SMX complex, which mediates recombination intermediate resolution^25,44^.

We previously showed that the SLX4-interacting protein SLX4IP, a scaffolding protein, also limits BLM-dependent recombination intermediate dissolution at ALT telomeres, acting in a pathway parallel to SLX4^23^. Loss of SLX4IP causes a BLM-dependent hyper-ALT phenotype characterized by increased APB formation, elevated levels of extrachromosomal telomeric circles, and accumulation of catenated telomere aggregates in mitosis. While our previous work established a role for SLX4IP in balancing BLM-dependent recombination intermediate dissolution, whether it also regulates other aspects of BLM activity at ALT telomeres remained unclear.

Here, we uncover a novel role for SLX4IP in regulating BLM-driven replication stress at ALT telomeres. Loss of SLX4IP increases telomeric replication stress and ATR signalling in ALT-positive cells. Strikingly, combined loss of SLX4IP and FANCM results in synthetic lethality that is rescued by BLM depletion, indicating that SLX4IP and FANCM act in parallel to restrain BLM. Mechanistically, we find that SLX4IP deficiency reduces replication fork dynamics and exacerbates BLM-dependent lagging-strand replication stress. Together, our findings identify SLX4IP as a key negative regulator of ALT telomere maintenance that limits BLM-driven replication stress.

## RESULTS

### SLX4IP depletion causes ATR-dependent replication stress at ALT telomeres

In ALT-positive cells, loss of SLX4IP induces a hyper-ALT phenotype driven by excessive BLM activity at telomere recombination intermediates^23^. Given the broad roles of BLM in ALT maintenance, regulating both replication stress and recombination, we speculated that the hyper-ALT phenotype imparted by SLX4IP depletion is not solely due to defects in recombination intermediate processing, but also arises from misregulation of ALT-dependent replication stress that occurs upstream of telomere recombination. To investigate this, we assessed the effect of SLX4IP depletion in ALT-positive cells on telomere fragility, replication dynamics and the activation of ATR-dependent signalling at telomeres, which are indicators of replicative stress^22,45^. First, we observed a significant increase in telomeric fragility in SLX4IP knockout U2OS cells (Figure 1A-B and Supplementary Figure 1A), indicating increased levels of replicative stress. This was further supported by reduced EdU incorporation in SLX4IP-deficient cells (Figure 1C-D), indicating impaired DNA synthesis and inefficient replication progression. Consistent with this, DNA combing analysis revealed that SLX4IP depletion in U2OS cells led to a decrease in replication fork speed compared to wild-type cells (Figure 1E). This effect was further exacerbated upon addition of 50 μM of the replication stress-inducing agent hydroxyurea (HU) (Figure 1E), further supporting a role for SLXIP in mitigating replication stress. Consistent with this, we also detected elevated levels of the ATR-dependent replication stress markers pSer33-RPA (Figure 1F-G) and pS345-CHK1 (Figure 1H-I) at telomeres. Notably, we observed increased telomeric localization of SLX4IP following 24h treatment with 4mM HU (Supplementary Figure 1B-C), suggesting that SLX4IP may be actively recruited to ALT telomeres under replication stress conditions. To determine whether this stress-induced recruitment is restricted to telomeres, we conducted CUT&Tag profiling of SLX4IP in both ALT-positive U2OS cells and ALT-negative HEK293 cells, a technique considerably more sensitive than microscopy-based approaches. CUT&Tag allowed us to detect SLX4IP accumulation at additional genomic regions including promoters (9.91% in U2OS, 6.83% in HEK293), gene bodies (23.94% in U2OS, 20.03% in HEK293), distal intergenic regions (37.65% in U2OS, 40.41% in HEK293), and fragile sites (23.24% in U2OS, 25.05% in HEK293) (Supplementary Figure 1D). Importantly, upon HU treatment, SLX4IP occupancy increased at all sites except promoters (Supplementary Figure 1E-G), indicating a potentially broader role in responding to replication stress genome-wide.

**Figure 1.**
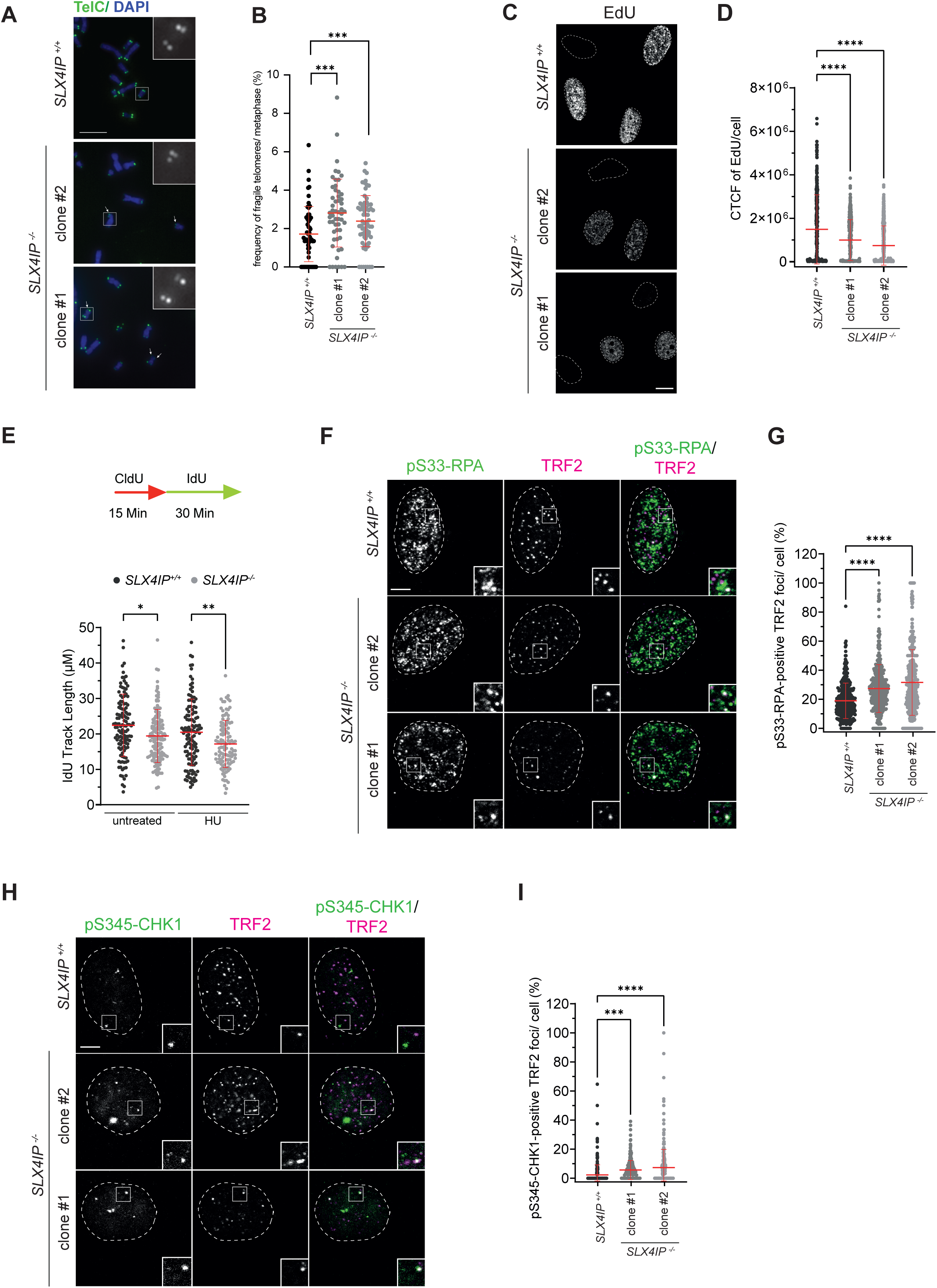
SLX4IP depletion activates the replication stress response at ALT telomeres. (A) U2OS cells were fixed, and metaphases were processed for telomere PNA (TelC) FISH and DAPI. Insets are 3X magnifications of the indicated fields. Scale bar represents 100 μm. Arrows indicate fragile telomeres. (B) Quantification of (A). The number of fragile telomeres counted in each metaphase was normalized to metaphases size. Data are represented as mean ± SD; n=2 with at least 30 metaphases per experiment; ***p < 0.001, one-way ANOVA. (C) U2OS cells were subjected to a 30 min EdU pulse before fixation, followed by staining of EdU via a Click-IT reaction. Dotted line represents DAPI (not shown). Scale bar represents 10 μm. (D) Quantification of (B). The EdU signal was quantified as corrected total nuclear intensity per cell. At least 100 cells per condition and experiment were counted. Data are represented as mean ± SD; n=3; ****p < 0.0001, one-way ANOVA. (E) U2OS cells were labelled with CldU and IdU and treated with 50 μM Hydroxyurea for the indicated times before DNA combing. Individual IdU fiber lengths are plotted. Median is indicated. *P* values were derived from ANOVA with Dunn’s multiple comparisons post-test. (F) U2OS cells were pre-extracted, fixed and processed for pSer33-RPA and TRF2 immunofluorescence. Insets are 3X magnifications of the indicated fields. Dotted line represents DAPI (not shown). Scale bar represents 5 μm. (G) Quantification of (E). The percentages of TelC foci that overlap with pSer33-RPA foci in each cell in were quantified. At least 100 cells per condition and experiment were counted. Data are represented as mean ± SD; n=3; ****p < 0.0001, one-way ANOVA. (H) U2OS cells were pre-extracted, fixed and processed for pSer345-CHK1 and TRF2 immunofluorescence. Insets are 3X magnifications of the indicated fields. Dotted line represents DAPI (not shown). Scale bar represents 5 μm. (I) Quantification of (G). The percentages of TelC foci that overlap with pSer345- CHK1 foci in each cell were quantified. At least 100 cells per condition and experiment were counted. Data are represented as mean ± SD; n=2; ****p < 0.0001, one-way ANOVA.

Together, these results support a role for SLX4IP in mitigating replication stress at ALT telomeres and possibly other genomic loci, ensuring efficient fork progression and preventing excessive activation of ATR-dependent replication stress signalling.

### SLX4IP acts in parallel to FANCM to limit replication stress at ALT telomeres

Since SLX4IP depletion induces replication stress at ALT telomeres, we sought to determine whether additional factors contribute to this process. FANCM, a translocase that limits replication stress at ALT telomeres^18,19,22,24^, emerged as a strong candidate because its loss shares several striking similarities with SLX4IP depletion. Specifically, both SLX4IP and FANCM loss leads to elevated replication stress at ALT telomeres and a BLM-dependent hyper-ALT phenotype^18,23^. In addition, both proteins interact directly with BLM, although the mechanistic significance of these interactions with regards to BLM activity remain unclear^23,46^. These common phenotypes suggested two possible scenarios: SLX4IP and FANCM could function in the same pathway or act in parallel pathways to counteract replication stress at ALT telomeres. To distinguish between these possibilities, we examined the phenotypic consequences of their combined depletion. Strikingly, simultaneous loss of SLX4IP and FANCM led to a significant augmentation of the replication stress markers pSer33-RPA and pCHK1 at ALT telomeres (Figure 2A-D and Supplementary Figure 2A-B), a further increase in APB formation (Figure 2E-F and Supplementary Figure 2C), and a pronounced synthetic growth defect (Figure 2G-H and Supplementary Figure 2D) in ALT-positive U2OS cells. To determine whether the synthetic growth defect observed with FANCM was specific to this factor, we also examined SMARCAL1, a related DNA translocase that acts in a pathway parallel to FANCM, and that has also been implicated in ALT telomere maintenance^47,48^. However, co-depletion of SLX4IP and SMARCAL1 did not result in a synthetic growth defect, suggesting that SMARCAL1 and SLX4IP function epistatically at replicating ALT telomeres (Supplementary Figure 2E-G). In addition, we tested whether SLX4, a key interactor of SLX4IP, and its associated endonucleases, XPF and MUS81, exhibit synthetic lethality with FANCM in ALT-positive U2OS cells. Unlike SLX4IP, co-depletion of FANCM with either MUS81 or XPF did not reduce clonogenic survival (Supplementary Figure 2H-M). Co-depletion with SLX4 produced a small but consistent decrease in clonal cell growth. However, this effect was not statistically significant (Supplementary Figure 2H-J). These data suggest that the synthetic interaction with FANCM is specific to SLX4IP and is not shared by SLX4 or its associated endonucleases.

**Figure 2.**
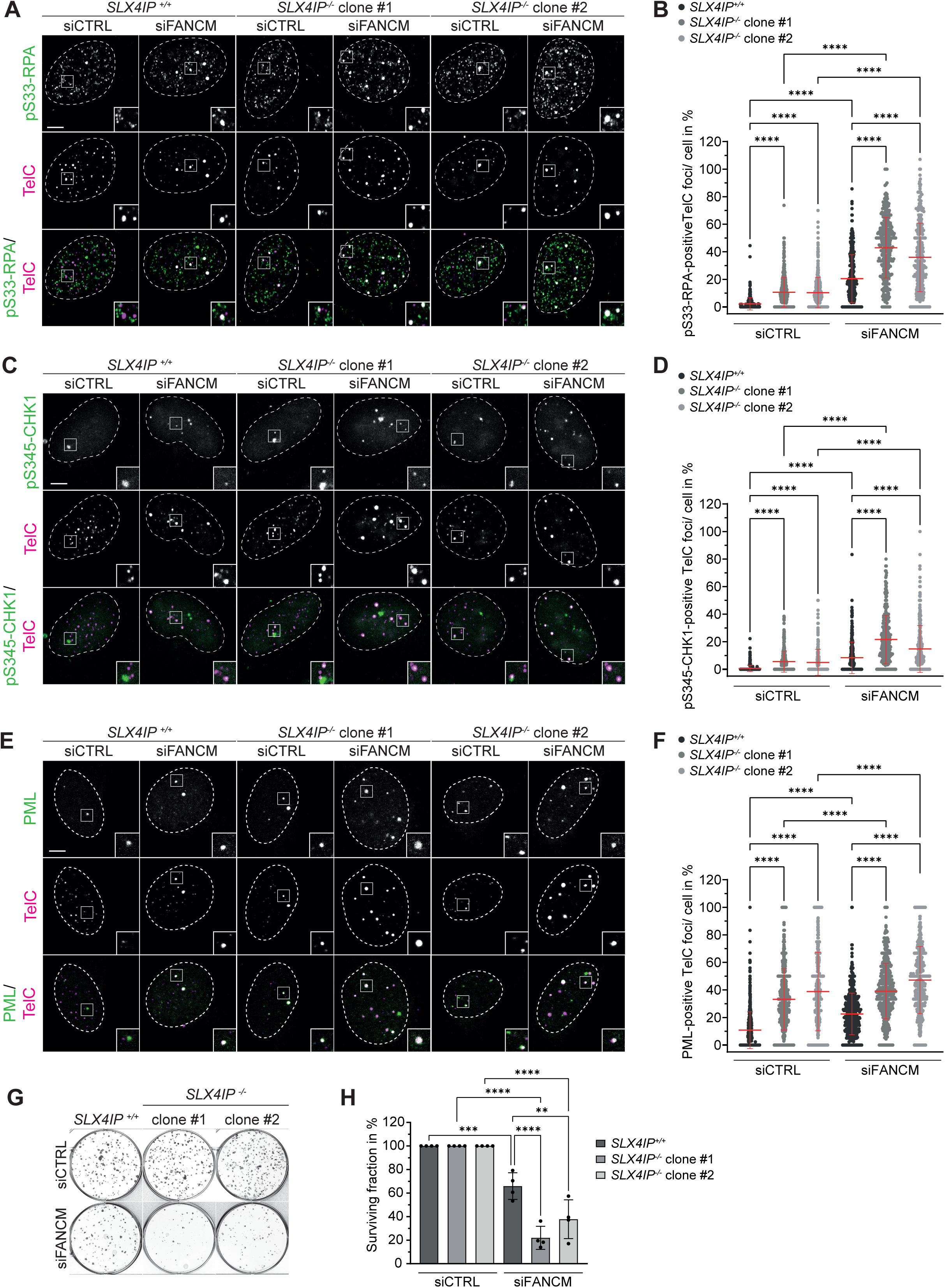
SLX4IP acts in parallel to FANCM to limit replication stress at ALT telomeres. (A) U2OS Cells were transfected either with non-targeting siRNA (siCTRL) or FANCM targeting siRNA (siFANCM), pre-extracted, fixed and processed for pSer33-RPA immunofluorescence followed by telomeric PNA (TelC) FISH. Insets are 3X magnifications of the indicated fields. Dotted line represents DAPI (not shown). Scale bar represents 5 μm. (B) Quantification of (A). The percentages of TelC foci that overlap with pSer33-RPA foci in each cell were quantified. At least 100 cells per condition and experiment were counted. Data are represented as mean ± SD; n=3; ****p < 0.0001, one-way ANOVA. Knockdown validations are shown in Supplementary Figure 2A. (C) U2OS cells were transfected either with non-targeting siRNA (siCTRL) or FANCM targeting siRNA (siFANCM), pre-extracted, fixed and processed for pSer345- CHK1 immunofluorescence followed by telomeric PNA (TelC) FISH. Insets are 3X magnifications of the indicated fields. Dotted line represents DAPI (not shown). Scale bar represents 5 μm. (D) Quantification of (C). The percentages of TelC foci that overlap with pSer345- CHK1 foci in each cell were quantified. At least 80 cells per condition and experiment were counted. Data are represented as mean ± SD; n=3; ****p < 0.0001, one-way ANOVA. Knockdown validations are shown in Supplementary Figure 2B. (E) U2OS cells were transfected either with non-targeting siRNA (siCTRL) or FANCM targeting siRNA (siFANCM), pre-extracted, fixed and processed for PML immunofluorescence followed by telomeric PNA (TelC) FISH. Insets are 3X magnifications of the indicated fields. Dotted line represents DAPI (not shown). Scale bar represents 5 μm. (F) Quantification of (E). APBs were quantified as the percentage of TelC foci that overlap with PML foci in each cell. At least 95 cells per condition and experiment were counted. Data are represented as mean ± SD; n=5; ****p < 0.0001, one-way ANOVA. Knockdown validations are shown in Supplementary Figure 2C. (G) U2OS cells were transfected either with non-targeting siRNA (siCTRL) or FANCM targeting siRNA (siFANCM) and seeded in a clonogenic survival assay. (H) Quantification of (G). Data are represented as mean ± SD; n=4; ****p < 0.0001, ***p < 0.001, **p < 0.01, one-way ANOVA. Knockdown validations are shown in Supplementary Figure 2D.

Together, our findings indicate that SLX4IP and FANCM function in distinct but complementary pathways to counteract replication stress at ALT telomeres.

### The synthetic lethal interaction between SLX4IP and FANCM is dependent on BLM

ALT-telomeric replication stress induced by FANCM is dependent on BLM activity^18^. Given the non-epistatic genetic relationship of SLX4IP and FANCM, we hypothesized that the increased levels of replication stress observed in SLX4IP-deficient cells may also be mediated by BLM. To test this possibility, we first examined the telomeric accumulation of BLM at SLX4IP-deficient ALT telomeres. Consistent with previous findings^23^, we observed significantly elevated total levels of BLM in SLX4IP-deficient whole-cell extracts (Figure 3A). Importantly, the increase in BLM levels led to an enhanced association of BLM with telomeres, a phenotype that was further exacerbated when FANCM was co-depleted (Figure 3B-C and Supplementary Figure 3A). Given this extensive telomeric accumulation of BLM, we next investigated whether the increased replication stress seen in SLX4IP-depleted cells, as well as the synthetic growth defect observed upon simultaneous depletion of SLX4IP and FANCM, might be dependent on BLM activity. To test this, we co-depleted BLM in SLX4IP-deficient cells as well as in SLX4IP/FANCM-co-depleted cells and assessed the impact on pSer33-RPA, cell viability and APB formation. Strikingly, loss of BLM rescued the elevated pSer33-RPA signals observed at SLX4IP-deficient cells, indicating a restoration of normal replication dynamics at ALT telomeres (Figure 3D-E and Supplementary Figure 3B). Furthermore, co-depletion of BLM rescued the synthetic growth defect observed upon simultaneous loss of SLX4IP and FANCM, demonstrating that the cytotoxicity that results from combined SLX4IP and FANCM deficiency is dependent on excessive BLM activity (Figure 3F-G and Supplementary Figure 3C). In addition, the increase in APB formation observed in SLX4IP/FANCM co-depleted cells was significantly reduced upon BLM loss (Supplementary Figure 3A,D-E), further supporting a model in which hyperactive BLM causes a hyper-ALT phenotype through excessive replicative stress in the absence of SLX4IP and FANCM.

**Figure 3.**
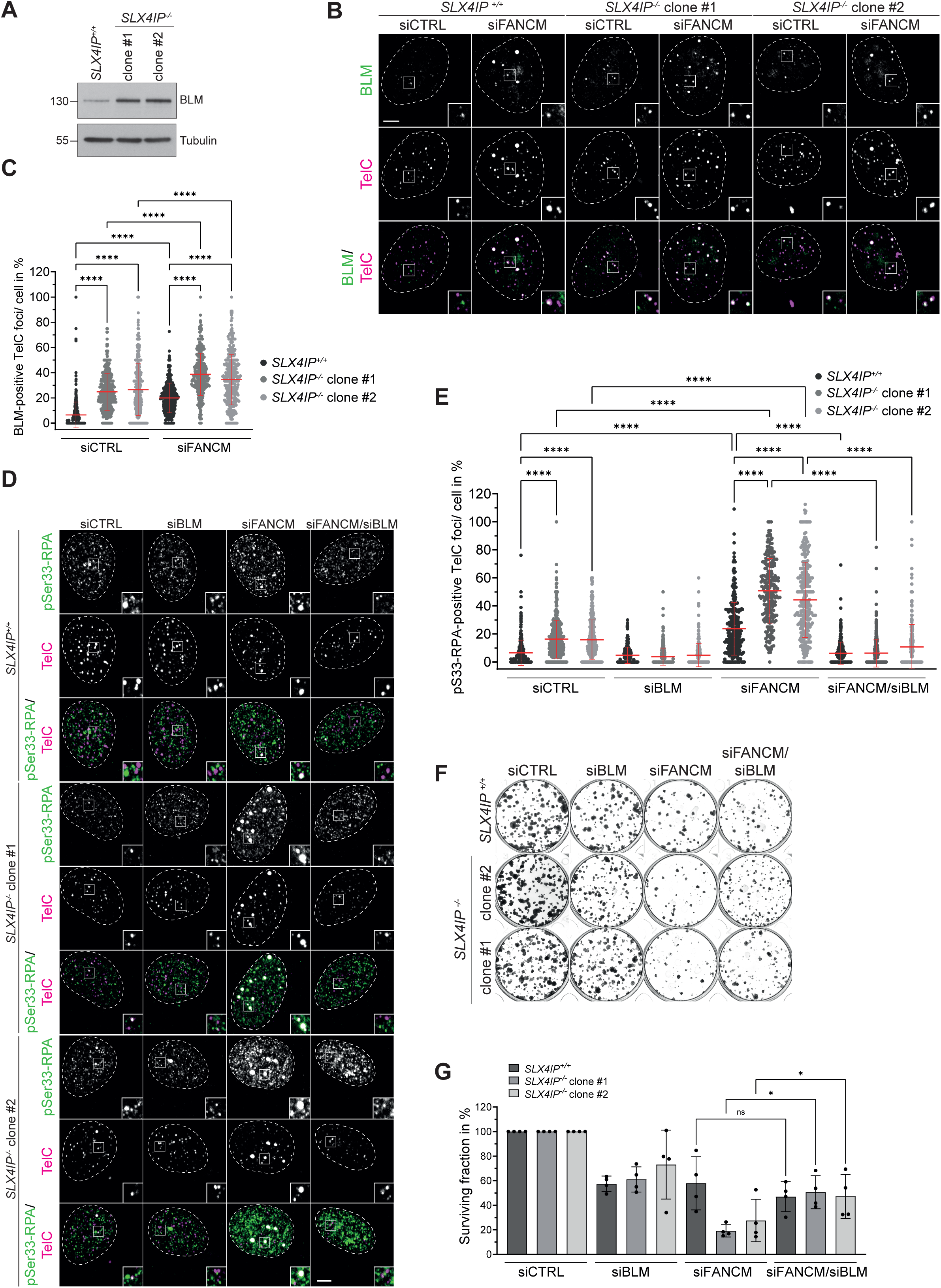
The synthetic lethal interaction between SLX4IP and FANCM is dependent on BLM. (A) Whole cell lysates of U2OS cells were separated by SDS-PAGE and analyzed for BLM levels by immunoblotting. Tubulin was used as loading control. The numbers on the left denote the molecular weight in kDa. (B) U2OS cells were transfected either with non-targeting siRNA (siCTRL) or FANCM targeting siRNA (siFANCM), pre-extracted, fixed and processed for BLM immunofluorescence followed by telomeric PNA (TelC) FISH. Insets are 3X magnifications of the indicated fields. Dotted line represents DAPI (not shown). Scale bar represents 5 μm. (C) Quantification of (B). The percentages of TelC foci that overlap with BLM foci in each cell were quantified. At least 95 cells per condition and experiment were counted. Data are represented as mean ± SD; n=3; ****p < 0.0001, one-way ANOVA. Knockdown validations are shown in Supplementary Figure 3A. (D) U2OS cells were transfected either with non-targeting siRNA (siCTRL) or knocked down for FANCM (siFANCM), BLM (siBLM) or both (siFANCM/siBLM). Cells were pre-extracted, fixed and processed for pSer33-RPA immunofluorescence followed by telomeric PNA (TelC) FISH. Insets are 3X magnifications of the indicated fields. Dotted line represents DAPI (not shown). Scale bar represents 5 μm. (E) Quantification of (D). The percentages of TelC foci that overlap with pSer33-RPA foci in each cell were quantified. At least 68 cells per condition and experiment were counted. Data are represented as mean ± SD; n=3; ****p < 0.0001, one-way ANOVA. Knockdown validations are shown in Supplementary Figure 3B. (F) U2OS cells were transfected either with non-targeting siRNA (siCTRL) or knocked down for FANCM (siFANCM), BLM (siBLM) or both (siFANCM/siBLM). Cells were then seeded in a clonogenic survival assay. (G) Quantification of (F). Data are represented as mean ± SD; n=4; *p < 0.05, one-way ANOVA; ns, not significant. Knockdown validations are shown in Supplementary Figure 3C.

Together, these findings indicate that SLX4IP depletion leads to a BLM-dependent hyper-replication stress phenotype at ALT telomeres. This suggests that SLX4IP functions as a critical regulator of BLM activity to maintain telomere stability and preserve cellular viability in ALT-positive cells, not only at the level of the telomere recombination intermediate^23^, but also to prevent excessive replication stress.

### SLX4IP depletion causes BLM-dependent lagging-strand replication stress

To further dissect the nature of the replication stress in SLX4IP knockout cells, we investigated whether this stress arises from defects in replication fork stability or an alternative mechanism. FANCM promotes replication fork remodelling at stalled forks and is synthetic lethal with the SMARCAL1 translocase^47,49^. Unlike FANCM, SLX4IP depletion did not result in synthetic lethality with SMARCAL1 (Supplementary Figure 2E-G), suggesting that the replication stress in SLX4IP-deficient cells does not stem from defects in fork remodelling.

To gain further mechanistic insight, we performed iPOND (isolation of proteins on nascent DNA) analysis to assess replication fork composition in SLX4IP-deficient U2OS cells^50^. Notably, we observed increased deposition of core histones as well MacroH2A1, H2AZ and H1 variants at replication forks (Figure 4A). These changes in histone dynamics are consistent with replication stress and suggest an altered fork environment in the absence of SLX4IP^51–53^. Additionally, SLX4IP depletion resulted in a threefold enrichment of the flap endonuclease FEN1, a key enzyme involved in Okazaki fragment processing, linking the observed replication stress to lagging-strand DNA synthesis. Consistent with this, clonogenic survival assays revealed that SLX4IP is epistatic with DNA Ligase 1, the enzyme responsible for sealing Okazaki fragment in clonogenic survival assays, suggesting that SLX4IP acts in the same pathway to ensure lagging-strand maturation (Supplementary Figure 4A-C). To assess the presence of single-stranded gaps in replicating DNA, which often arise from defects in Okazaki fragment maturation, we performed DNA combing combined with S1 nuclease treatment. S1 nuclease selectively cleaves single-stranded regions, thereby shortening DNA fibres that contain replication-associated gaps. Compared to untreated controls, S1-treated samples exhibited a significant reduction in replication tract length, which was further decreased in SLX4IP-deficient cells (Figure 4B). This increased susceptibility to S1 digestion suggests indicates a higher burden of single-stranded gaps in the absence of SLX4IP, consistent with impaired lagging strand maturation or incomplete gap filling. Supporting this, we observed a significant increase in nuclear poly(ADP-ribose) (PAR) signal, which predominantly marks unligated Okazaki fragments in unstressed cells (Figure 4C-D)^54,55^. In addition, we detected elevated levels of SLX4IP on chromatin following treatment with 1 μ or 10 μ doses of the PARP inhibitor Olaparib, suggesting a role for SLX4IP in managing replication-associated PARylation or unligated Okazaki fragments (Supplementary Figure 4D-G). Finally, given that BLM was recently shown to unwind immature Okazaki fragments at ALT telomeres^20^, we next tested whether the lagging-strand replication stress observed in SLX4IP-deficient cells was dependent on BLM activity. To this end, we co-depleted BLM in SLX4IP knockout cells and again assessed PAR levels as a proxy for lagging-strand replication stress. Remarkably, BLM depletion rescued the elevated PAR signal in SLX4IP-deficient cells back to wildtype levels (Figure 4E-F and Supplementary Figure 4H), which indicates a restoration of baseline replication dynamics and stress.

**Figure 4.**
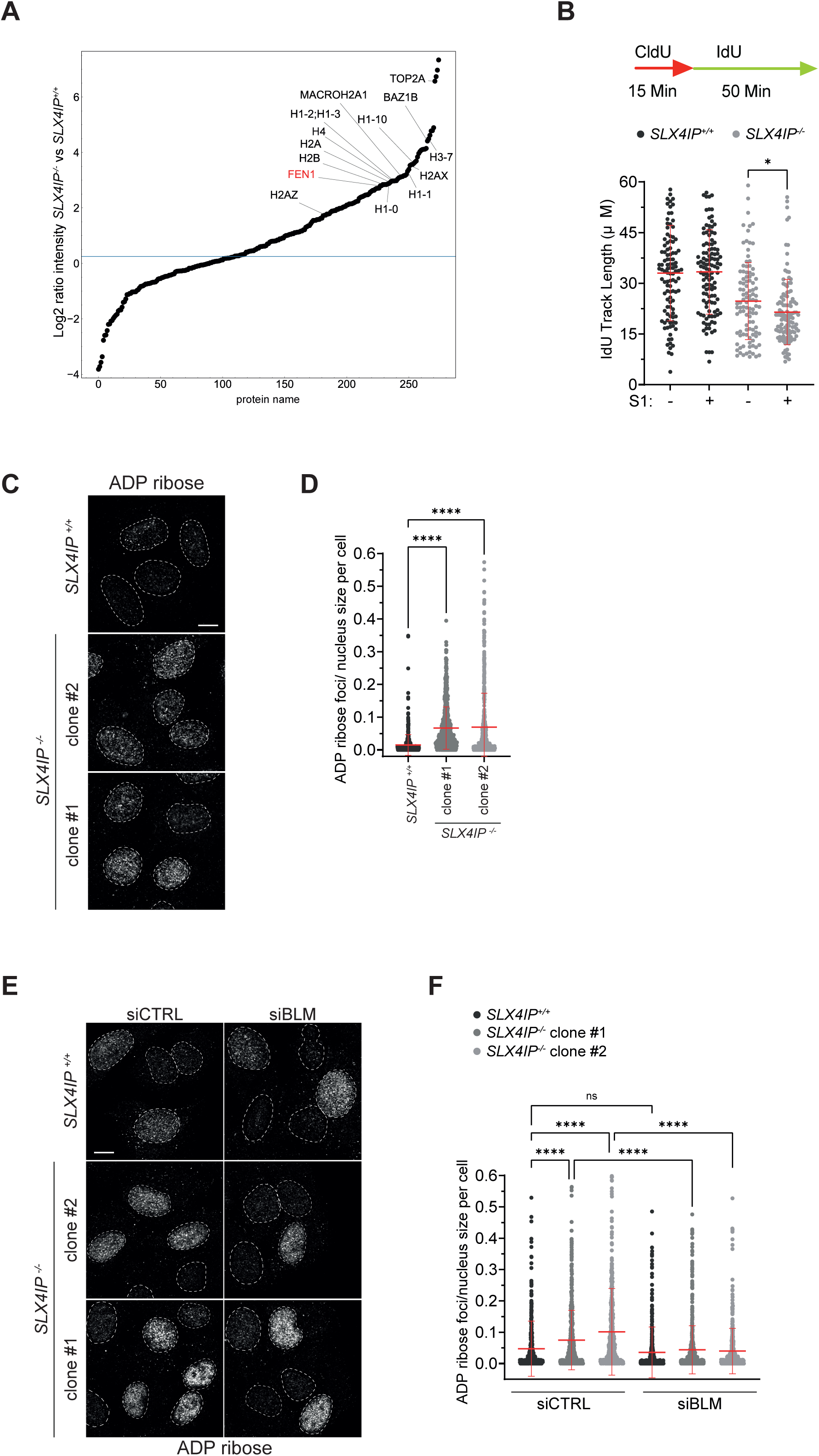
SLX4IP depletion causes BLM-dependent lagging-strand replication stress. (A) Loss of SLX4IP accumulates FEN1 and histones at replication forks. U2OS were subjected to a 20 min EdU pulse to label newly synthesized DNA, followed by iPOND coupled to mass spectrometry. The graph shows the log2 ratio intensity of proteins accumulated at replication forks in *SLX4IP^-/-^* compared to *SLX4IP^+/+^* cells. (B) U2OS cells were labelled with CldU and IdU before DNA combing and in gel S1 nuclease treatment. Individual IdU fibre lengths are plotted. Median is indicated. *P* values were derived from ANOVA with Dunn’s multiple comparisons post-test. (C) U2OS cells were treated with 10 µM PARG inhibitor for 30 min, immediately fixed and processed for ADP ribose immunofluorescence. Dotted line represents DAPI (not shown). Scale bar represents 10 μm. (D) Quantification of (C). ADP ribose foci in each cell were quantified and normalized to the nucleus size. At least 100 cells per condition and experiment were counted. Data are represented as mean ± SD; n=3; ****p < 0.0001, one-way ANOVA. (E) U2OS cells were transfected either with non-targeting siRNA (siCTRL) or BLM targeting siRNA (siBLM). Cells were treated with 10 µM PARG inhibitor for 30 min, immediately fixed and processed for ADP ribose immunofluorescence. Dotted line represents DAPI (not shown). Scale bar represents 10 μm. (F) Quantification of (E). ADP ribose foci in each cell were quantified and normalized to the nucleus size. At least 100 cells per condition and experiment were counted. Data are represented as mean ± SD; n=3; ****p < 0.0001, one-way ANOVA; ns, not significant. Knockdown validations are shown in Supplementary Figure 4H.

Together, these findings suggest that SLX4IP functions to suppress aberrant BLM activity at ALT telomeres and prevent excessive unwinding of unligated Okazaki fragments^20^. In the absence of SLX4IP, hyperactive BLM disrupts lagging-strand processing, increasing replication stress and compromising telomere stability.

## DISCUSSION

Here, we characterize SLX4IP as a key regulator of BLM helicase activity during ALT telomere replication, revealing a previously unrecognized role in mitigating replication stress associated with lagging-strand DNA synthesis. We demonstrate that depletion of SLX4IP causes BLM-dependent telomeric replication stress that is marked by increased ATR signalling and impaired replication fork dynamics. Additionally, we identify a synthetic lethal interaction between SLX4IP and the FANCM translocase, suggesting that these two proteins act in parallel pathways to mitigate BLM-dependent replication stress at ALT telomeres. While our data do not directly visualize lagging-strand replication intermediates, the enrichment of FEN1 at SLX4IP-deficient replication forks and the increase in nuclear PAR, combined with the BLM dependency, support a model in which SLX4IP limits lagging-strand replication stress by constraining the BLM-mediated unwinding of unligated Okazaki fragments. This mechanism may be especially critical for preserving telomere stability in ALT-positive cells.

ALT telomeres are particularly prone to replication challenges, in part due to BLM-mediated unwinding of immature Okazaki fragments^20,56^. Our findings position SLX4IP as a modulator of this process. In the absence of SLX4IP, unchecked BLM activity disrupts lagging-strand maturation, generating 5’flaps and single-stranded DNA gaps that trigger PARP1/2-dependent PARylation^20,57^, activate ATR signalling, and promote hyperrecombination. By constraining BLM activity, SLX4IP prevents this cytotoxic cascade and helps preserve telomere integrity.

Both replication stress and recombination must be finely balanced in ALT cells to ensure telomere stability^6^. Our results underscore that while ALT cells upregulate BLM helicase to manage the replication stress and recombination intermediates generated by the ALT process, this upregulation is not without consequences as demonstrated by the aberrant unwinding of immature Okazaki fragments on the lagging strand^20^. Mechanisms to regulate BLM activity are therefore essential. While FANCM has previously been implicated in counteracting BLM-dependent replication stress^18^, our findings suggest that SLX4IP plays a complementary role in limiting replication stress in ALT cells. Together, SLX4IP and FANCM act to counteract the hyperactivity of BLM, ensuring that replication stress is managed without compromising genomic stability and inducing non-productive hyperrecombination. The upregulation of SLX4IP in ALT-positive cells^23^, suggests an adaptive response to heightened BLM activity in these cells. This adaptive upregulation helps to keep BLM-dependent replication stress and recombination within a tolerable range that supports telomere maintenance while preventing excessive genomic instability.

An outstanding question remains regarding the precise mechanism by which SLX4IP counteracts BLM activity. Although SLX4IP directly interacts with BLM, it does not directly affect its helicase activity in vitro^23^. Whether SLX4IP modulates BLM helicase activity in vivo remains unclear. One possibility is that SLX4IP, which lacks enzymatic activity, serves as a scaffolding factor that regulates the large BLM interactome in a context-dependent manner^58^. Alternatively, and not mutually exclusively, SLX4IP may act downstream of hyperactive BLM to mitigate its effects, preventing excessive replication stress and hyperrecombination.

Beyond its role at ALT telomeres, the ability of SLX4IP to limit replication stress suggests that it may also function at other genomic loci. This idea is supported by our CUT&Tag analysis, which revealed that SLX4IP accumulates at non-telomeric regions, particularly following HU-induced replication stress. Importantly, we observed this chromatin distribution in both ALT and non-ALT cells, hinting at broader functions for SLX4IP beyond its role at ALT telomeres. Certain regions, such as common fragile sites, repetitive sequences, and highly transcribed loci, experience elevated levels of replication stress due to their structural and chromatin features^59^. Notably, SLX4IP was previously been shown to be enriched at common fragile sites alongside the BTR complex^60^, raising the possibility that it contributes to the maintenance of such vulnerable regions. While BLM is involved in replication fork remodelling throughout the genome, SLX4IP may have a more restricted role in regulating replication stress at specific loci, rather than serving as a global modulator of BLM activity.

ALT cells are particularly vulnerable to FANCM loss^18,19,22,24^, which supports the idea of FANCM inhibition as a therapeutic strategy for ALT cancers^61^. Notably, SLX4IP is inactivated in a subset of ALT cancers^23^. The ALT-specific synthetic lethal interaction between SLX4IP and FANCM suggests that the efficacy of FANCM inhibitors may be enhanced in SLX4IP-deficient ALT tumours. Therefore, SLX4IP could serve as a biomarker to predict the effectiveness of FANCM-targeted therapies in ALT-positive cancers.

In summary, this work defines a previously unrecognized role for SLX4IP in ALT telomere maintenance, where it acts to restrain BLM activity during lagging-strand DNA synthesis. Our data underscores the importance of precise regulation of replication stress and recombination in ALT-positive cells, ensuring productive telomere elongation while preventing excessive genomic instability. Moreover, our findings open new avenues for investigating SLX4IP as a potential regulator of BLM activity at non-telomeric loci and as a biomarker for therapeutic strategies targeting ALT-associated replication stress.

## MATERIALS AND METHODS

### Cell culture

U2OS and HEK293 cells were cultured in Dulbecco’s modified eagle medium with glutamax (Life Technologies) supplemented with 10% (v/v) fetal bovine serum (FBS). Cells were maintained using standard tissue culture procedures and kept in a humidified incubator set to 37°C and 5% CO2. Cells were regularly tested for Mycoplasma. Cells were frozen in 90% (v/v) FBS and 10% (v/v) DMS0, transferred to Mr. Frosty freezing containers (Nalgene) and kept for at least 1 day at -80 °C. For long-term storage, cells were transferred into a liquid N2 tank.

### RNA interference

RNAi transfections were performed using DharmaFECT 1 (Horizon) in a forward transfection mode. U2OS cells were treated with a final amount of 40 nM siRNA. The transfection reagent volumes used were according to the manufacturer’s manual. Cells were plated to obtain 30-40 % confluency on the day of transfection. Next day, Dharmafect 1 and Opti-MEM were mixed by vortexing, shortly spun down and incubated for 5 min at RT. siRNA was added to the transfection mix, vortexed, spun down and incubated for 20 min at RT. Meanwhile, cell culture medium of the cells was replaced with fresh medium. After incubation, medium was added to the transfection reagent/siRNA complex, briefly vortexed, shortly spun down and dispensed dropwise onto the cells. 5-7 h after transfection, the medium was substituted for fresh medium. Cells were further processed 48- 72 h post-transfection.

### Indirect immunofluorescence

Cells were grown on #1.5 glass coverslips in a 24-well plate. Where indicated, cells were treated with hydroxyurea (HU; 4 mM, 24 h) and water as control or with Olaparib (1 µM or 10 µM, 48 h) and DMSO as control before fixation. In siRNA knockdown experiments, cells were fixed approximately 72 h post transfection. Cells were washed with 1 mL/well 1X PBS once at RT and pre-extracted with 0.5 mL/well ice-cold pre- extraction buffer (20 mM HEPES pH 7.5, 20 mM NaCl, 5 mM MgCl2, 300 mM sucrose, 0.5 % NP-40, 1 mM DTT, phosphatase and protease inhibitor tablets) for 20 min on ice. The pre-extraction buffer was removed and cells were immediately fixed with 2 % (w/v) formaldehyde (Life Technologies) in PBS for 20 min at RT. After fixation, cells were washed with 1X PBS three times for 5 min respectively and then blocked with 0.5 mL/well Antibody Dilution Buffer (ADB; 10 % (v/v) normal goat serum, 0.1 % (v/v) Triton X-100, 0.1 % (v/v) saponin in PBS) for at least 1 h. Cells were incubated with 30 µL primary antibody diluted in ADB in a humid chamber for 1 h at RT. The primary antibodies were diluted as follows: mouse anti-SLX4IP 1:50, rabbit anti-BLM 1:1000, mouse anti-PML 1:1000, rabbit anti-pS33-RPA 1:1000, rabbit anti-pS345-CHK1 1:50 and mouse anti-TRF2 1:500. After incubation in primary antibodies, cells were washed once with 1X PBS + 0.1 % (v/v) Triton-X-100 for 5 min, followed by two washes with 1X PBS for 5 min. Each coverslip was then counterstained with 300 µL Alexa Fluor secondary antibodies (Thermo Fisher Scientific) diluted 1:1000 in ADB, for 1 h at RT. After washing once with 1X PBS + 0.1 % (v/v) Triton-X-100 for 5 min, cells were washed with 1X PBS two times for 5 min. To visualize nuclei, cells were stained with 1 mL/well 0.1 µg/mL DAPI in 1X PBS for 5 min, followed by two more washes with 1 mL/well 1X PBS. Coverslips were mounted onto glass slides with 10 µL drops of Prolong Diamond Antifade mounting agent (Life Technologies) and left to dry overnight at RT in the dark. Slides were stored at 4 °C. Pre-extraction, fixation, blocking, all wash steps, the secondary antibody incubation and DAPI staining were carried out on a horizontal shaker. Starting from the addition of the fluorescent secondary antibodies, coverslips were always incubated in the dark.

For immunofluorescence of ADP ribose, cells were grown on #1.5 glass coverslips in a 24-well plate. Before fixation, live cells were treated with 10 µM PARG inhibitor PDD00017273 (Selleckchem) for 30 min. They were washed with 1 mL/well 1X PBS once at RT and fixed with 0.5 mL/well ice-cold 100% methanol for 15 min at -20°C. Methanol was kept at -20°C overnight before using it for fixation. After fixation cells were washed with 1 mL/well 1X PBS for three times for 5 min each. Cells were always processed the same day for ADP ribose immunofluorescence. For that, cells were blocked with 0.5 mL/well blocking buffer (1X PBS / 5 % (v/v) normal goat serum / 0.3 % (v/v) Triton X-100) for 1 h at RT. Cells were then incubated in 300 µL rabbit anti-Poly/Mono-ADP Ribose (Cell Signaling) diluted 1:3200 in antibody dilution buffer (1X PBS / 1 % (w/v) BSA / 0.3 % (v/v) Triton X-100) overnight at 4°C. After washing once with 1X PBS + 0.1 % (v/v) Triton-X-100 for 5 min, cells were washed with 1X PBS two times for 5 min. Each coverslip was then counterstained with 300 µL goat anti-rabbit Alexa Fluor 488 secondary antibody (Thermo Fisher Scientific) diluted 1:1000 in ADB, for 1 h at RT. After washing once with 1X PBS + 0.1 % (v/v) Triton-X-100 for 5 min, cells were washed with 1X PBS two times for 5 min. To visualize nuclei, cells were stained with 1 mL/well 0.1 µg/mL DAPI in 1X PBS for 5 min, followed by two more washes with 1 mL/well 1X PBS. Coverslips were mounted onto glass slides with 10 µL drops of Prolong Diamond Antifade mounting agent (Life Technologies) and left to dry overnight at RT in the dark. Slides were stored at 4 °C. All antibody incubations, all wash steps and DAPI stainings were carried out on a horizontal shaker. Starting from the addition of the fluorescent secondary antibody, coverslips were always incubated in the dark.

### Telomeric Peptide Nucleic Acid Fluorescence In Situ Hybridization (PNA-FISH)

Cells were treated with 0.2 µg/ml of colcemid for 90 minutes to arrest cells in metaphase. Trypsinized cells were then incubated in 75 mM KCl for 20 min and pelleted at 1000rpm for 5 min, fixed with methanol:acetic acid (3:1), spread on glass slides and left overnight at room temperature to dry. The slides were rehydrated in PBS for 5 minutes, fixed in 4% formaldehyde for 5 minutes, treated with 1 mg/ml of pepsin for 10 minutes at 37°C, and fixed in 4% formaldehyde for 5 minutes. Next, slides were dehydrated in 70%, 85%, and 100% (v/v) ethanol for 15 minutes each and then air-dried. Metaphase chromosome spreads were hybridized with a telomeric TAMRA- TelG 5’-(TTAGGG)3-3’ PNA probe (Bio-synthesis) in hybridizing solution (70% formamide, 0.5% blocking reagent (Roche), 10mM Tris-HCl pH 7.2) for 90 seconds at 80°C followed by 2 hours at room temperature and washed twice with washing buffer (70% formamide, 10mM Tris-HCl pH 7.2) for 15 min at room temperature. Slides were mounted using ProLong Gold antifade with DAPI (Life Technologies).

### Immunofluorescence coupled to fluorescence in situ hybridization (IF-FISH)

Samples were processed as described in the section ‘indirect immunofluorescence’. After incubating cells with Alexa Fluor secondary antibodies (Thermo Fisher Scientific), cells were washed once with 1 mL/well 1X PBS + 0.1 % (v/v) Triton-X- 100 for 5 min followed by two washes with 1 mL/well 1X PBS for 5 min. After that, cells were fixed again with 1 mL/well 2 % (w/v) formaldehyde in PBS for 20 min at RT and then washed three times with 1 mL/well 1X PBS. Next, coverslips were dehydrated on ice in 0.5 mL/well ice-cold 70 %, 95 %, and 100 % (v/v) ethanol for 5 minutes each and then air-dried. Coverslips were processed no later than the next day after dehydration. Glass slides were preheated for 1 min on a heat block set to 75°C. Then 30 µL drops of hybridizing solution (10 nM TelC-Cy5 PNA probe (PNA Bio) in 70 % (v/v) formamide, 0.5 % (w/v) blocking reagent (Roche), 10 mM Tris-HCl pH 7.2) were placed on the glass slides and heated for 1 min. Dry coverslips were placed with cells facing downwards into the hybridizing solution and hybridized for 90 seconds at 75 °C followed by 2 h at RT in a humid chamber. After incubation, cells were washed twice with 0.5 mL/well washing buffer (70 % (v/v) formamide, 10 mM Tris-HCl pH 7.2) for 15 min at RT. After that, coverslips were washed with 1 mL/well 1X PBS for 5 min. To visualize nuclei, cells were stained with 1 mL/well 0.1 µg/mL DAPI in 1X PBS for 5 min, followed by two more washes with 1 mL/well 1X PBS. Coverslips were mounted onto glass slides with 10 µL drops of Prolong Diamond Antifade mounting agent (Life Technologies). Secondary antibody incubations, all wash steps and DAPI stainings were carried out on a horizontal shaker. Starting from the addition of the fluorescent secondary antibodies, coverslips were always incubated in the dark.

### EdU Click-IT reaction

Cells were grown on #1.5 glass coverslips in a 24-well plate. Before fixation, live cells were incubated in 0.5 mL/ well medium supplied with 10 µM EdU for 30 min. They were washed with 1 mL/well 1X PBS once at RT and pre-extracted with 0.5 mL/well ice-cold pre-extraction buffer (0.2 % (v/v) Triton-X-100 in 1X PBS) for 10 min on ice. The pre-extraction buffer was removed and cells were immediately fixed with 4 % (w/v) formaldehyde (Life Technologies) in PBS for 15 min at RT. The EdU incorporation was visualized with the Click-iT Plus EdU Alexa Fluor 488 Imaging Kit (Thermofisher). Permeabilisation, blocking, washing steps and the EdU Click-IT reaction were performed according to the manufacturer’s manual. To each coverslip 250 µL of the Click-iT Plus reaction cocktail was added. After washing cells with 1 mL/well of 3% (w/v) BSA in 1X PBS, cells were washed two more times with 1X PBS for 5 min. To visualize nuclei, cells were stained with 1 mL/well 0.1 µg/mL DAPI in 1X PBS for 5 min, followed by two more washes with 1 mL/well 1X PBS. Coverslips were mounted onto glass slides with 10 µL drops of Prolong Diamond Antifade mounting agent (Life Technologies). Blocking, the EdU Click-IT reaction, all wash steps and the DAPI staining were carried out on a horizontal shaker.

### Image acquisition and analysis of IF data

Images of IF and IF-FISH stainings were acquired with a Leica SP8-X or SP8-DLS inverted confocal microscope equipped with a 63X glycerol objective. Following acquisition, images were imported into ImageJ (NIH) for automated quantification.

Automated quantification of foci, colocalization analysis of foci and nuclear size measurements were performed with a custom script in Fiji. To correct foci numbers to the nuclear size, the number of foci of each nucleus was divided by the respective nucleus area. For the corrected total nuclear fluorescence intensity (CTNF), the integrated density and nuclear sizes were quantified automatically with a custom Fiji script. Background measurements were taken manually from each image by choosing 3 regions of interest in each image and calculating their average mean intensity. The CTNF was then calculated by the following equation: Integrated density – (Area of nucleus * average mean intensity of background).

### Whole-cell extracts

Cells were rinsed with 1X PBS, trypsinized and collected in DMEM. Cells were pelleted by centrifugation at 300 g for 5 min and washed with 1X PBS. Cell pellets were snap-frozen on dry ice and stored at -80 °C. For lysis, cell pellets were thawed on ice and resuspended in lysis buffer (50 mM Hepes-KOH, pH 7.5, 100 mM KCl, 2 mM EDTA, 0.5 % IGEPAL, 10 % glycerol, 1 mM DTT, 1X protease and phosphatase inhibitors (Pierce Protease Inhibitor Tablets EDTA-free, Pierce Phosphatase Inhibitor Mini Tablets, Thermo Fisher Scientific)). Lysates were incubated on ice for 30 min and sonicated in the bioruptur for 10 cycles 30 s on and off. Cell lysates were clarified by centrifugation at 13 000 g for 20 min at 4 °C. Protein concentration was determined using the BCA method (DC protein assay (Biorad)) according to the manufacturer’s instructions. Lysates were denatured in 1X Laemmli Sample Buffer and 5 mM DTT for 5 min at 95-100 °C and stored at -20 °C.

### SDS-PAGE and immunoblotting

Proteins were separated by SDS-PAGE. Proteins were transferred onto a nitrocellulose membrane (Biorad) at 100 V for 2 h. After transfer, the membrane was blocked in 5 % (w/v) skim milk/ TBST (1X TBS/ 0.1 % (v/v) Tween-20) for at least 30 min at RT and incubated with the indicated primary antibody diluted in 5% skim milk/ TBST for 1 h at RT or overnight at 4nC. The primary antibodies sheep anti-SLX4IP, mouse anti-Vinculin, rabbit anti-BLM and mouse anti-alpha-Tubulin were diluted 1:1000. Rabbit anti-FANCJ was diluted 1:300 in TBST. The membrane was then washed 5 times for 5 min with TBST, incubated with a horseradish peroxidase-conjugated secondary antibody diluted 1:5000 in 5% skim milk/ TBST for 1 h at RT, and washed again 5 times for 5 min with TBST. The immunoblot was developed on Hyperfilm (Fisher Scientific) using ECL Western Blotting Reagent (Thermo Fisher Scientific). All incubations were carried out on a horizontal shaker.

### Clonogenic survival assay

Around 48 h after siRNA transfection, cells were trypsinized, strained with 40 µM cell strainers, counted and re-plated into 6-well dishes in conditioned medium. For that, conditioned medium was collected from confluent cell culture flasks and filtered with a sterile 0.22 µM filter. Generally, conditioned medium was stored up to 2 weeks at 4 °C. Each condition was plated into a 6-well plate in duplicates or triplicates. Remaining cells were collected in cell pellets for validating knockdowns by qPCR. For the survival assay, cells were grown for 8-11 days and fixed in a 20% (v/v) methanol/0.4% (w/v) crystal violet solution for 10 min. Crystal violet was removed and residual dye was washed off with deionized water. Plates were placed upside down and left to dry overnight. Plates were imaged with the GelCount scanner (Oxford Optronix) in a 600 dpi resolution and images were saved as tif files. The area percentage of colonies covering each well was analysed with a custom Fiji script.

### RT-qPCR

Cells were centrifuged at 300 g for 5 min. Medium was discarded and cells were resuspended in 1 mL DPBS and transferred into RNAse-free microcentrifuge tubes. Cells were centrifuged again at 500 g for 5 min and the supernatant was discarded. Cell pellets were washed one more time with 0.5 mL DPBS, snap-frozen on dry ice and stored at -80°C. RNA isolation was performed with the RNeasy Mini Kit (Qiagen) according to the manufacturer’s instructions. RNA was reverse transcribe into cDNA with the Luna Script RT SuperMix Kit (Biolabs). RT-qPCR was performed using Luna Universal qPCR Master Mix-SYBR green (NEB).

### iPOND coupled to mass spectrometry

#### iPOND

U2OS cells were labeled with 10mM EdU for 20 minutes. Cells were cross- linked in 1% fresh formaldehyde/PBS for 10 minutes (RT) and quenched with 1.25 M glycine. Cells were permeabilized in 0.25% Triton-X/PBS and washed once with 0.5% BSA/PBS followed by PBS. Click reaction was performed for 1 hr with PEG4-biotin azide (Invitrogen). The cells were subsequently lysed by sonication using a Diagenode Pico sonicator on ultra-high for 30 minutes. DNA-protein complexes were purified using streptavidin C1 magnetic beads for 1 hr. Samples were washed (5 minutes each) with lysis buffer (1% SDS in 50 mM Tris, pH 8.0), low salt buffer (1% Triton X-100, 20 mM Tris, pH 8.0, 2 mM EDTA, pH 8.0, 150 mM NaCl), high salt buffer(1% Triton X-100, 20 mM Tris, pH 8.0, 2 mM EDTA, pH 8.0, 500 mM NaCl), lithium chloride buffer (100 mM Tris, pH 8.0, 500 mM LiCl, 1% Igepal), followed by two washes in lysis buffer. Captured proteins were eluted and cross-links were reversed in 2X SDS sample buffer for 30 min at 95 C.

### In-Gel Digestion for mass spectrometry

Unless otherwise specified, all procedures were carried out at room temperature. Samples were separated using 4–12% Tris-glycine mini gels (Novex™, Invitrogen). Following electrophoresis, gels were fixed for 30 minutes in a solution containing 50% methanol and 1.2% phosphoric acid to immobilize proteins. After a brief rinse with water, gels were stained overnight in a solution composed of 50% methanol, 1.2% phosphoric acid, 1.3 M ammonium sulfate, and 0.1% (w/v) Coomassie Brilliant Blue G-250. Each lane was excised using a gel cutting tool and divided into two fractions: fraction 2, containing the streptavidin band, and fraction 1, comprising the remaining protein content. Each fraction was cut into small cubes, transferred into microcentrifuge tubes, and subjected to multiple rounds of destaining and washing to remove residual dye and salts. The washing procedure consisted of sequential 15-minute incubations: first in 50 mM ammonium bicarbonate, followed by a second incubation in 50 mM ammonium bicarbonate with 50% ethanol. After each incubation, the supernatant was discarded. For heavily stained gel pieces, incubations were carried out at 60 °C. This washing cycle was repeated 2–3 times until gel pieces were completely destained. Prior to reduction and alkylation, gel pieces were dehydrated in 100% ethanol for 10 minutes and briefly dried in a vacuum concentrator to remove residual liquid. Reduction of cysteine residues was carried out by incubating gel pieces in 20 mM DTT, 50 mM ammonium bicarbonate for 20 minutes. After removal of the reducing solution, alkylation was performed using 80 mM chloroacetamide in 50 mM ammonium bicarbonate for 20 minutes. Excess reagents were eliminated through a series of three 15-minute washes: (i) 50 mM ammonium bicarbonate, (ii) 25 mM ammonium bicarbonate with 50% ethanol, and (iii) 100% ethanol. Gel pieces were then dried completely using a vacuum concentrator. Tryptic digestion was performed overnight at 37 °C by adding 0.25 µg of trypsin (Promega) in 50 µL of 50 mM ammonium bicarbonate to each sample fraction. Peptides were extracted the next day by two 15-minute elutions in 150 µL of extraction buffer (80% acetonitrile, 0.2% formic acid) under sonication. For desalting and final cleanup, peptide samples were reconstituted in 200 µL of 0.2% formic acid and processed via solid-phase extraction (SPE) using C18 cartridges (1 cc, 50 mg sorbent, Sep-Pak, Waters) connected to a vacuum manifold. Cartridges were equilibrated with 100% acetonitrile and washed twice with 0.2% formic acid. After sample loading, two additional washes with 0.2% formic acid were performed. Peptides were then eluted using 80% acetonitrile with 0.2% formic acid and dried completely by vacuum centrifugation. Prior to LC–MS analysis, samples were reconstituted in 0.2% formic acid and quantified using a NanoDrop spectrophotometer.

### LC-MS and data processing

Liquid chromatography–mass spectrometry (LC–MS) analysis was performed using a Vanquish Neo UHPLC system (Thermo Fisher Scientific) coupled to an Orbitrap Astral mass spectrometer (Thermo Fisher Scientific) via a Nanospray Flex ion source. Peptide separation was carried out on a 40 cm in-house–packed fused silica C18 column, maintained at 50 °C using a custom-built column heater. The spray voltage was set to 2 kV, and the ion transfer tube temperature was held at 280 °C. For each run, 500 ng of peptide from streptavidin-containing fractions was injected, while 1 µg was loaded for all other samples. Chromatographic separation was achieved at a flow rate of 300 nL/min using a 30-minute linear gradient from 5% to 46% solvent B (solvent A: 0.2% formic acid in water; solvent B: 80% acetonitrile with 0.2% formic acid). Data acquisition was performed on the Orbitrap Astral mass spectrometer with the following settings. MS1 spectra were acquired in the Orbitrap at a resolution of 240,000 over a scan range of 300–1350 m/z, using a maximum injection time of 50 ms, a normalized automatic gain control (AGC) target of 250%, and an RF lens setting of 30%. Data-dependent acquisition (DDA) included the following filters: (i) precursor charge states from +2 to +6 were selected; (ii) monoisotopic peak selection (MIPS) was enabled and set to “peptide”; and (iii) dynamic exclusion was applied with a 6-second exclusion window and a ±3 ppm mass tolerance. Selected precursors were fragmented using higher-energy collisional dissociation (HCD) with a normalized collision energy of 26%. MS2 spectra were acquired in the Astral analyzer with an isolation window of 0.8 m/z, a scan range of 150–2000 m/z, a maximum injection time of 5 ms, a normalized AGC target of 300%, and an RF lens setting of 50%. The DDA cycle time was set to 0.6 seconds. RAW files generated from the LC–MS runs were analyzed with MaxQuant (version 2.6.2.0), searching against a UniProt human FASTA file containing both SwissProt and TrEMBL entries. Gel fractions originating from the same lane were assigned to identical experiment names but treated as separate fractions in MaxQuant. With the few exceptions mentioned below, all MaxQuant parameters were set to their default values. In the ”Group-specific parameters” tab, “Label-free quantification” was enabled. In the Global parameters tab, Match between runs enabled. Raw files generated from LC–MS analyses were processed using MaxQuant (version 2.6.2.0). Spectra were searched against a UniProt human FASTA database (downloaded February 2024) containing both SwissProt and TrEMBL entries. Gel fractions originating from the same lane were assigned identical experiment names but were treated as separate fractions within MaxQuant. In the “Group-specific parameters” tab, label-free quantification (LFQ) was enabled. Additionally, the “Match between runs” feature was activated under the global parameters. All other MaxQuant parameters were kept at default values. For downstream data analysis, the proteinGroups.txt output file generated by MaxQuant was used. Label-free quantification (LFQ) intensity values from replicates 1 and 2 of both wild-type (WT) and SLX4IP-knockout (SLX4IP-KO) samples were log2- transformed. For each replicate pair, a SLX4IP-KO to WT log2 ratio was subsequently calculated. Proteins identified by LC-MS/MS are listed in Supplementary Table 1.

### DNA Combing

Cells were labeled with 20 μM CldU (5-chloro-2’-deoxyuridine) followed by 100 μM IdU (5-Iodo-2’-deoxyuridine), for the time schemes indicated, with or without 50 μM Hydroxyurea (Sigma-Aldrich, 4000465) or 50 uM Cisplatin. Approximately 600,000 cells were embedded in 1.5% low-melting agarose in phosphate-buffered saline (PBS) and digested overnight in 0.1% sarkosyl, proteinase K (2 mg/ml), and 50 mM EDTA (pH 8.0) at 50°C. Agarose plugs were washed in TE (10mM Tris pH=8.0, 1mM EDTA) transferred to 100 mM MES (pH 5.7), melted at 68°C, and digested with 1.5 U of β- agarase overnight at 42°C. DNA was combed onto silanized coverslips (Automate Scientific) using a custom combing apparatus (Erasmus MC, Rotterdam). The DNA was stained with antibodies that react with IdU and CldU for 1 hour, washed in PBS, and probed with secondary antibodies for 45 min. For S1 Nuclease assay, cells were treated as in the DNA molecular combing assay above. Plugs were washed once and switched to 1× S1 nuclease buffer. Plugs were switched to fresh S1 nuclease buffer, and S1 nuclease (10 U) was added. Plugs were incubated for 1 hour at 37°C. The reaction was immediately quenched with 50 mM EDTA. Plugs were washed in TE (pH 8.0) once with 50 mM EDTA and then transferred to 100 mM MES (pH 5.7) before continuing the DNA molecular combing protocol as described above. All images were obtained using a 40× oil objective (Nikon Eclipse Ti2, Kinetix Full-frame Camera). Analysis of fiber lengths was performed using Nikon Elements software.

### CUT&Tag

#### Protein A-Tn5 purification and assembly of transposomes

The 3XFLAG tag was removed from the 3Xflag-pA-Tn5-Fl plasmid (Addgene, #124601) through molecular cloning. For this, the 3Xflag-pA-Tn5-Fl plasmid was amplified using mutagenesis primers 5’-ACCATGGGTATGACCATGATTACGCC-3’ and 5’- CATGGTCATACCCATGGTATATCTCCTTC-3’, and subsequently cloned using the In-Fusion HDCloning Plus (Takara Bio, #638911). Protein A-Tn5 was then expressed and purified as previously described^62^. Annealing of Tn5MEDS was performed by mixing equal amounts of 200µM pMENTS oligo with 200 µM Tn5ME-A or 200 µM Tn5ME-B followed by incubation at 95°C for 2 minutes. The temperature was subsequently decreased to 25°C in 5°C increments, with each step held for 5 minutes. Annealed Tn5MEDS were stored at -20°C. For transposome assembly, each 4 µL of 200 µM annealed Tn5MEDS-A and Tn5MEDS-B were combined with 100 µL of the protein A-Tn5 glycerol stock and incubated for 50 minutes at 23°C. Assembled transposomes were stored at -20°C.

#### Cleavage Under Targets and Tagmentation (CUT&Tag)

For nuclei isolation, HEK293 or U2OS cells treated with or without 4mM hydroxyurea (HU) for 24h were incubated in NE1 buffer (20 mM HEPES-KOH pH 7.9, 10 mM KCl, 0.5 mM Spermidine, 0.1 % Triton X-100, 20 % glycerol, 1X cOmplete Proteinase Inhibitor) on ice for 5 min or 20 min, respectively. Nuclei were fixed with 0.1% formaldehyde (Thermo Scientific, 28906) for 2 minutes at RT followed by quenching with 75 mM glycine. Nuclei were pelleted by centrifugation at 1,300*xg* for 4 min at 4°C and washed with PBS. CUT&Tag experiments were performed as described previously^62^, with minor modifications: Nuclei were not immobilized on concanavalin A-conjugated beads. Instead, nuclei were pelleted to change supernatant through centrifugation at 600*xg* for 4 minutes at RT. All wash buffers were supplemented with 0.01% digitonin and 0.01% NP-40. Briefly, 1x10^5^ fixed nuclei were washed with 150-wash buffer (20 mM HEPES pH 7.5, 150 mM NaCl, 0.5 mM spermidine, Protease inhibitor, 0.01% digitonin, 0.01% NP-40) and incubated with mouse anti-SLX4IP (sc-377066, Santa Cruz Biotechnology) 1:100 in 50 µl of primary antibody binding buffer (150-wash buffer, 1% BSA, 2 mM EDTA) at 4° overnight. The next day, nuclei were incubated rabbit anti-mouse antibody (1:100; ab46540, abcam) for 1h at RT, followed by a wash with 150-wash buffer. Nuclei were resuspended in 50 µl of 300-wash buffer (20 mM HEPES pH7.5, 300 mM NaCl, 0.5 mM spermidine, Protease inhibitor, 0.01% digitonin, 0.01% NP-40) with 1:200 dilution of pA-Tn5 P7/P5 adapter complex and incubated for 1 hour at RT rotating. Nuclei were washed with 300-wash buffer followed by tagmentation in 50 µl of tagmentation buffer (300-wash buffer, 10 mM MgCl2). Tagmented DNA was isolated using DNA Clean & Concentrator kit (Zymo Research, #D4004) directly after tagmentation and eluted from the column in 23 µl Nuclease-free H2O. Libraries were amplified with barcoded i7 and i5 primers with NEBNext High- Fidelity 2X PCR Master Mix (NEB, M0541L) for 17 cycles. Optimal number of amplification cycles was determined via qPCR. Amplified libraries were subjected to double-sided size selection using 0.67x and 1.3x of High-Prep PCR magnetic beads (Magbio, #AC-60050). Libraries were sequenced for 100 cycles paired-ended on NextSeq1000 platforms. CUT&Tag experiments were performed with two biological per cell line with each two technical replicates.

### CUT&Tag Bioinformatics

#### Generation of a non-redundant union peak set for SLX4IP CUT&Tag

For each cell line we combined the stringent SEACR peak calls obtained from all biological replicates (four ± HU and four ± SLX4IP, eight BED files in total per line) into a single, non-redundant region set that was used in all downstream analyses.

1. **Input.** Scaled, replicate-level peak BED files produced by SEACR were collected from the project directories
2. **Quality control.** A Bash wrapper halted with an error if no files were found and printed a warning if the expected eight files were not present, ensuring that every merge contained the full replicate set.
3. **Sorting.** All replicate peaks for a given cell line were concatenated and sorted with BEDTools sort (v 2.31.0; default parameters).
4. **Merging.** Overlapping or immediately adjacent intervals were collapsed with BEDTools merge (default parameters), generating the cell-line-specific union set.

#### Annotation of SLX4IP CUT&Tag peaks and generation of per-category summary

For each cell line we classified the union peak set into genomic categories and exported a tab-delimited summary table that was subsequently plotted in GraphPad Prism (10.4.2).

1. **Input:** The cell-line-specific union peak BED file was imported into R (v 4.3.0) with **GenomicRanges**(Bioconductor 3.18).
2. Annotation tracks:

a. **Gene models:** *TxDb.Hsapiens.UCSC.hg38.knownGene*. Promoters were defined as −1 kb/+250 bp around the transcription-start site; gene bodies comprised the remaining exonic + intronic sequence; distal regions were the 100 kb upstream flank exclusive of promoters.
b. **Fragile sites:** experimentally defined common fragile-site coordinates supplied as a custom BED file. All tracks were trimmed to standard chromosomes and harmonised to UCSC sequence-level style.
3. **Peak annotation and summary:** Full-containment overlaps were calculated with countOverlaps (default settings). Peaks were assigned in the priority order *FragileSite > Promoter > GeneBody > Distal*; peaks matching none of the tracks were labelled *Unannotated*. Category counts and percentages were written to per_category_peak_summary.tsv.

The same workflow was executed for HEK293 and U2OS, differing only in the input peak file.

#### Coverage quantification, RPKM normalisation and log₂(HU / SLX4IP) ratio calculation

For every genomic region set we quantified CUT&Tag signal in four merged libraries (U2OS HU, U2OS SLX4IP, HEK HU, HEK SLX4IP), converted the counts to RPKM and derived log₂ ratios between HU-treated and SLX4IP-depleted samples.

1. **Input.** Region BED files and four duplicate-filtered, hg38-aligned BAM files.
2. **Raw coverage.** Region-level read counts were extracted with BEDTools multicov (v 2.31.0, default parameters).
3. **Library-size normalisation.** Total mapped reads per BAM were obtained with samtools view -c (v 1.19). Counts were converted to

RPKM=reads in region×10^9^/region length (bp)×total mapped reads.

1. 4. **Ratio calculation.** For each region the log₂(HU / SLX4IP) ratio was computed separately for U2OS and HEK; a pseudocount of 1 × 10⁻⁶ was added to zero values before taking the logarithm.
2. 5. **Output.** Visualisation by GraphPad Prism (10.4.2).

### log₂(HU / SLX4IP) coverage tracks and heat-map visualisatio

For each cell line we generated log₂-ratio coverage tracks and visualised HU-induced changes across category-specific peak sets.

1. **Input.** CPM-normalised bigWig files for HU-treated and SLX4IP-depleted libraries (HEK293 and U2OS) together with BED files defining distal, fragile- site, promoter and gene-body peak subsets.
2. **Log₂-ratio bigWigs.** Coverage was compared with deepTools bigwigCompare (v 3.5.2; --operation log2 --pseudocount 1 -- skipZeroOverZero).
3. **Signal matrix.** Using computeMatrix reference-point (±2.5 kb around region centres, 50 bp bins, --missingDataAsZero), per-category matrices were created and exported as both compressed (*.gz) and tab-delimited text files.
4. **Visualisation.** Heat maps and average profiles were rendered with plotHeatmap (per-group mode, colour map *bwr*, zMin = -1.2, zMax = 1.2), yielding PDF figures that summarise log₂(HU / SLX4IP) signal distributions across the selected peak categories.

The workflow was executed independently for HEK293 and U2OS; only the input bigWigs and BED region files differed.

#### Genome browser visualisation of SLX4IP occupancy

IGV (v2.16.1) was used to generate genomic snapshots of a promoter and fragile site showing differential SLX4IP occupancy. Normalised coverages of SLX4IP pm) and annotated regional files were used as input for the visualisation.

## Materials

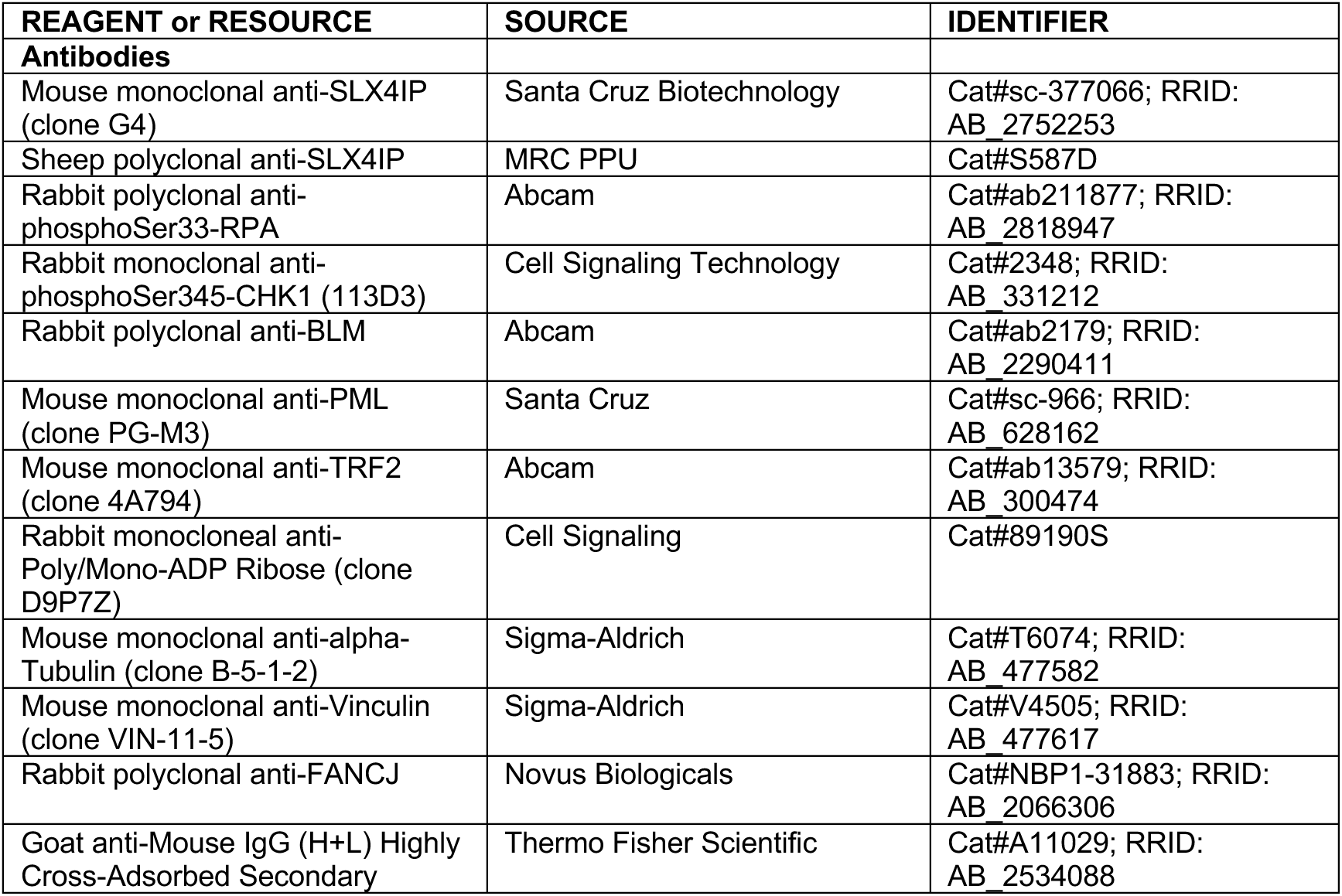
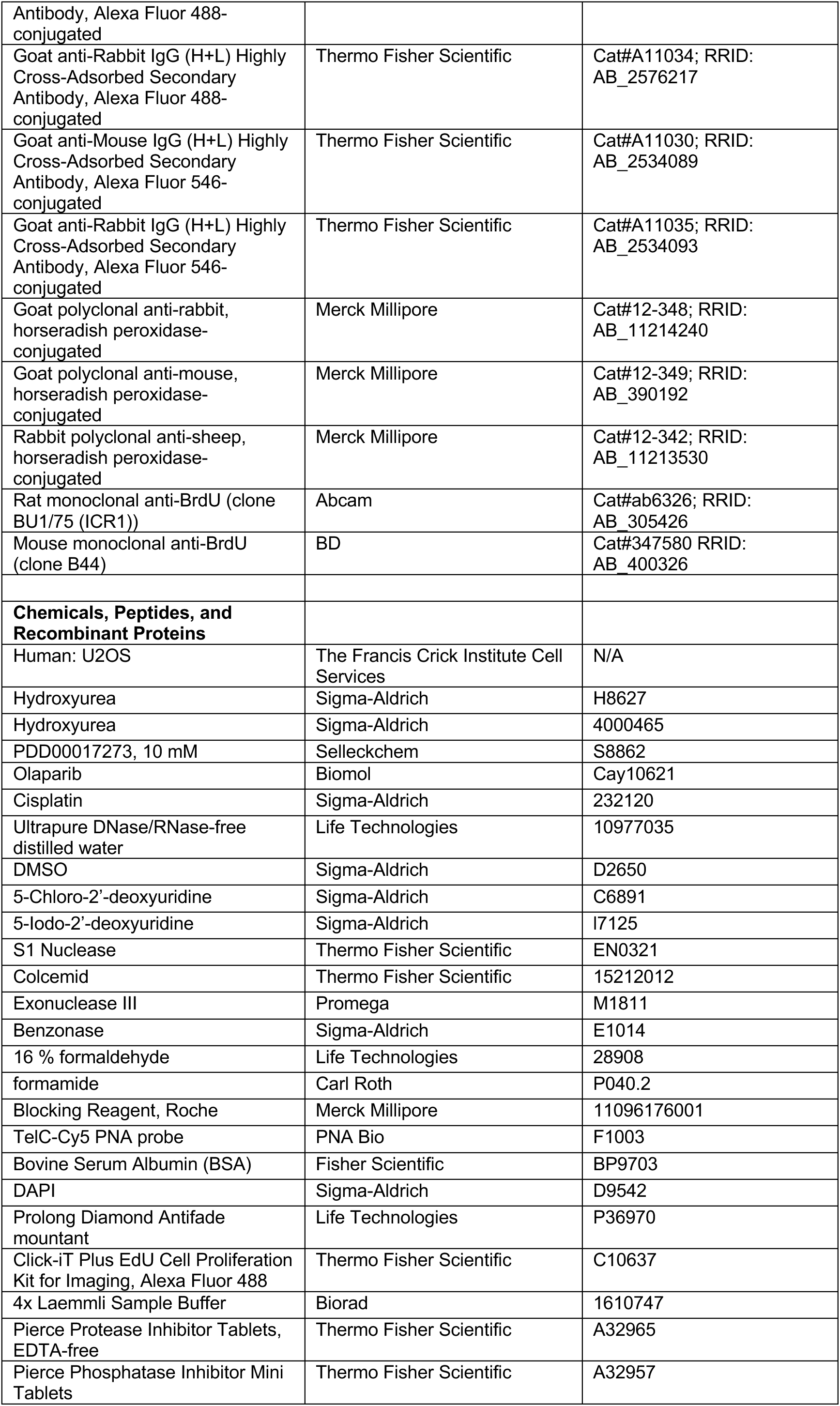
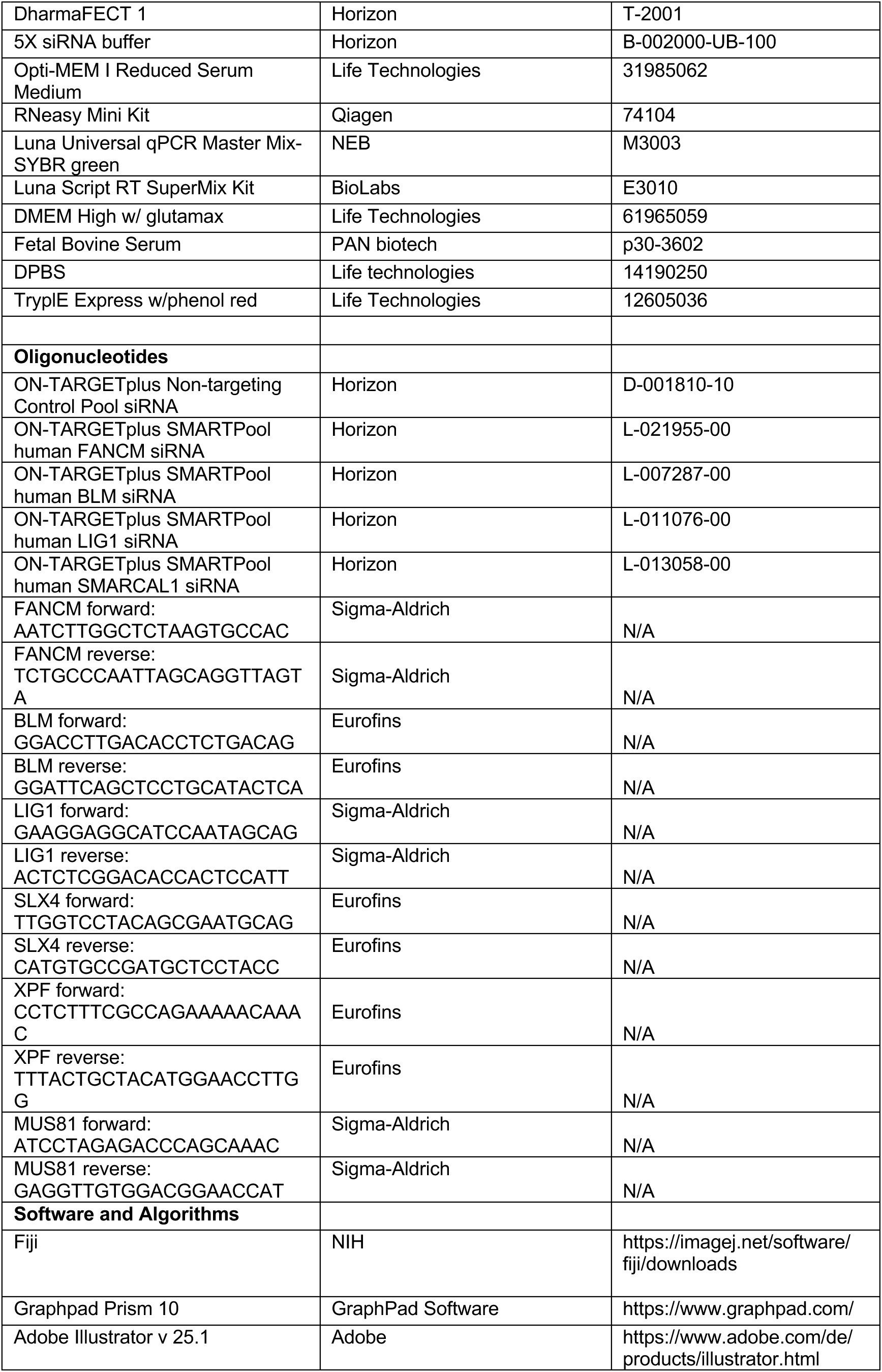

## ACKNOWLEDGMENTS

We acknowledge the assistance of Marine Duhamel and Jorge Bouças of the Bioinformatics Core Facility at the Max Planck Institute for Biology of Ageing in processing and analysis of the CUT&Tag data. Work in the J.C. lab is supported by NIH grant P41GM108538. Work in the K.P.M.M. lab is supported by NIH grant NIEHS R00ES034058; K.P.M.M was additionally supported by the Office of the Vice Chancellor for Research and Graduate Education, University of Wisconsin-Madison. This work was funded by the Deutsche Forschungsgemeinschaft (DFG, German Research Foundation) – FOR5504 – project number 496650118, granted to S.P.; work in the S.P. lab is additionally supported by core funding from the Max Planck Society. The funders had no role in study design, data collection and analysis, decision to publish, or preparation of the manuscript.

## AUTHOR CONTRIBUTIONS

S.P. and J.S. conceived the project and wrote the manuscript. S.P. performed the FISH experiment shown in Figure 1A; J.B. perfomed the immunoblotting experiment shown in Figure S1A; A.E.K. and P.P. performed the CUT&Tag experiment; R.H.H analysed the CUT&Tag data; S.B. performed all qPCR experiments; K.P.M.M performed the combing experiments and the iPOND pulldown; M.M. and J.C. performed the mass spectrometry anaylsis of the iPOND experiment; F.P. performed the experiments shown in Figures 3A, 4C-F and S2E-G. J.S. carried out all other experiments.

## SUPPLEMENTARY FIGURE LEGENDS

Supplementary Figure 1; related to Figure 1

(A) Whole cell lysates of U2OS cells were separated by SDS-PAGE and analyzed for SLX4IP levels by immunoblotting. Tubulin was used as loading control. The numbers on the left denote the molecular weight in kDa.

(B) U2OS cells were either treated with 4 mM hydroxyurea (HU) or water as control for 24 hours. Cells were then immediately pre-extracted, fixed and processed for SLX4IP immunofluorescence followed by telomeric PNA (TelC) FISH. Insets are 3X magnifications of the indicated fields. Dotted line represents DAPI (not shown). Scale bar represents 10 μm.

(C) Quantification of (B). SLX4IP foci in each cell were quantified and normalized to the nucleus size. At least 100 cells per condition and experiment were counted. Data are represented as mean ± SD; n=3; ****p < 0.0001, Student’s t-test.

(D) U2OS and HEK293 cells were processed for CUT&Tag analysis. Shown is the genomic annotation of SLX4IP ChIP-seq peaks in U2OS (upper panel) and HEK293 (lower panel) cells, displayed as the percentage of peaks falling within promoters, gene bodies, distal intergenic regions, and fragile sites.

(E) U2OS and HEK293 cells were treated with 4mM hydroxyurea (HU) for 24 h and processed for CUT&Tag analysis. Shown is the distribution of the log₂ ratio of normalized SLX4IP coverage (RPKM, reads per kilobase per million) after HU treatment versus untreated control. Each point represents one peak; a value > 0 denotes increased occupancy, < 0 denotes decreased occupancy. Red line in violin plot shows median ratio (log2).

(F) “Tornado” heat maps showing the HU-to-control coverage ratio for individual SLX4IP peaks, sorted within each genomic category in (D). Red indicates higher, blue lower SLX4IP signal after HU treatment. Related to Supplementary Figure 1E.

(G) Genome-browser snapshots of two representative fragile site (upper panel) and promoter (lower panel) loci. Shown is SLX4IP binding following HU exposure. Signal tracks are shown as log₂-normalized read coverage. Coverage is shown as counts per million (cpm). Related to Supplementary Figure 1E.

Supplementary Figure 2; related to Figure 2

(A) RNA was isolated from cell pellets of each condition, reverse transcribed into cDNA followed by RT-qPCR. Data were normalized to the siCTRL-treated samples; n=1. Related to Figure 2A-B.

(B) RNA was isolated from cell pellets of each condition, reverse transcribed into cDNA followed by RT-qPCR. Data were normalized to the siCTRL-treated samples; n=1. Related to Figure 2C-D.

(C) RNA was isolated from cell pellets of each condition, reverse transcribed into cDNA followed by RT-qPCR. Data were normalized to the siCTRL-treated samples. Data are represented as mean ± SD; n=5; ****p < 0.0001, one-way ANOVA. Related to Figure 2E-F.

(D) RNA was isolated from cell pellets of each condition, reverse transcribed into cDNA followed by RT-qPCR. Data were normalized to the siCTRL-treated samples. Data are represented as mean ± SD; n=4; ****p < 0.0001, one-way ANOVA. Related to Figure 2G-H.

(E) U2OS cells were transfected either with non-targeting siRNA (siCTRL) or siRNA targeting SMARCAL1 (siSMARCAL1). Cells were then seeded in a clonogenic survival assay.

(F) Quantification of (E). Data are represented as mean ± SD; n=3; one-way ANOVA; ns, not significant.

(G) RNA was isolated from cell pellets of each condition, reverse transcribed into cDNA followed by RT-qPCR. Data were normalized to the siCTRL-treated samples. Data are represented as mean ± SD; n=2; ****p < 0.0001, one-way ANOVA. Related to Supplementary Figure 2E-F.

(H) U2OS cells were transfected with the denoted combinations non-targeting siRNA (siCTRL), siFANCM, siSLX4 and siXPF. Cells were then seeded in a clonogenic survival assay.

(I) Quantification of (H). Data are represented as mean ± SD; n=3; one-way ANOVA; ns, not significant.

(J) RNA was isolated from cell pellets of each condition, reverse transcribed into cDNA followed by RT-qPCR. Data were normalized to the siCTRL-treated samples. Data are represented as mean ± SD; n=2; ****p < 0.0001, one-way ANOVA. Related to Supplementary Figure 2H-I.

(K) U2OS cells were transfected with the denoted combinations non-targeting siRNA (siCTRL), siFANCM and siMUS81. Cells were then seeded in a clonogenic survival assay.

(L) Quantification of (K). Data are represented as mean ± SD; n=3; one-way ANOVA; ns, not significant.

(M) RNA was isolated from cell pellets of each condition, reverse transcribed into cDNA followed by RT-qPCR. Data were normalized to the siCTRL-treated samples. Data are represented as mean ± SD; n=2; ****p < 0.0001, **p < 0.01, one-way ANOVA. Related to Supplementary Figure 2K-L.

Supplementary Figure 3; related to Figure 3

(A) RNA was isolated from cell pellets of each condition, reverse transcribed into cDNA followed by RT-qPCR. Data were normalized to the siCTRL-treated samples. Data are represented as mean ± SD; n=3; ****p < 0.0001, one-way ANOVA. Related to Figure 3C-D.

(B) RNA was isolated from cell pellets of each condition, reverse transcribed into cDNA followed by RT-qPCR. Data were normalized to the siCTRL-treated samples. Data are represented as mean ± SD; n=3; ****p < 0.0001, one-way ANOVA. Related to Figure 3C-D. Left panel: FANCM knockdown, right panel: BLM knockdown.

(C) RNA was isolated from cell pellets of each condition, reverse transcribed into cDNA followed by RT-qPCR. Data were normalized to the siCTRL-treated samples. Data are represented as mean ± SD; n=3; ****p < 0.0001, one-way ANOVA. Related to Figure 3E-F. Left panel: FANCM knockdown, right panel: BLM knockdown.

(D) U2OS cells were transfected either with non-targeting siRNA (siCTRL) or knocked down for FANCM (siFANCM), BLM (siBLM) or both (siFANCM/siBLM). Cells were pre-extracted, fixed and processed for PML immunofluorescence followed by telomeric PNA (TelC) FISH. Insets are 3X magnifications of the indicated fields. Dotted line represents DAPI (not shown). Scale bar represents 5 μm.

(E) Quantification of (D). APBs in were quantified as the percentage of TelC foci that overlap with PML foci in each cell. At least 68 cells per condition and experiment were counted. Data are represented as mean ± SD; n=3; ****p < 0.0001, one-way ANOVA. Knockdown validations are shown in Supplementary Figure 3A.

Supplementary Figure 4; related to Figure 4.

(A) U2OS cells were transfected either with non-targeting siRNA (siCTRL) or knocked down for Ligase1 (siLIG1), pre-extracted, fixed and processed for SLX4IP immunofluorescence. Insets are 3X magnifications of the indicated fields. Dotted line represents DAPI (not shown). Scale bar represents 5 μm.

(B) Quantification of (A). SLX4IP foci in each cell were quantified and normalized to the nucleus size. At least 100 cells per condition and experiment were counted. Data are represented as mean ± SD; n=3; ****p < 0.0001, Student’s t-test.

(C) RNA was isolated from cell pellets of each condition, reverse transcribed into cDNA followed by RT-qPCR. Data were normalized to the siCTRL-treated samples. Data are represented as mean ± SD; n=3; ****p < 0.0001, one-way ANOVA.

(D) U2OS cells were either treated with 1 µM Olaparib or DMSO as control for 48 hours. Cells were then immediately pre-extracted, fixed and processed for SLX4IP immunofluorescence. Insets are 3X magnifications of the indicated fields. Dotted line represents DAPI (not shown). Scale bar represents 5 μm.

(E) Quantification of (D). SLX4IP foci in each cell were quantified and normalized to the nucleus size. At least 100 cells per condition and experiment were counted. Data are represented as mean ± SD; n=2; ****p < 0.0001, Student’s t-test.

(F) U2OS cells were either treated with 10 µM Olaparib or DMSO as control for 48 hours. Cells were then immediately pre-extracted, fixed and processed for SLX4IP immunofluorescence. Insets are 3X magnifications of the indicated fields. Dotted line represents DAPI (not shown). Scale bar represents 5 μm.

(G) Quantification of (F). SLX4IP foci in each cell were quantified and normalized to the nucleus size. At least 80 cells per condition and experiment were counted. Data are represented as mean ± SD; n=3; ****p < 0.0001, Student’s t-test.

(H) RNA was isolated from cell pellets of each condition, reverse transcribed into cDNA followed by RT-qPCR. Data were normalized to the siCTRL-treated samples. Data are represented as mean ± SD; n=3; ****p < 0.0001, one-way ANOVA. Related to Figure 4E-F.

Supplementary Table 1; related to Figure 4

The detailed MS results of the iPOND experiment shown in Figure 4A.

## REFERENCES

1. Maciejowski, J. & De Lange, T. Telomeres in cancer: tumour suppression and genome instability. Nat Rev Mol Cell Biol 18, 175–186 (2017).

2. Bryan, T. M., Englezou, A., Dalla-Pozza, L., Dunham, M. A. & Reddel, R. R. Evidence for an alternative mechanism for maintaining telomere length in human tumors and tumor-derived cell lines. Nat Med 3, 1271–1274 (1997).

3. Henson, J. D. & Reddel, R. R. Assaying and investigating Alternative Lengthening of Telomeres activity in human cells and cancers. FEBS Letters 584, 3800–3811 (2010).

4. Dilley, R. L. & Greenberg, R. A. ALTernative Telomere Maintenance and Cancer. Trends in Cancer 1, 145–156 (2015).

5. Dilley, R. L. et al. Break-induced telomere synthesis underlies alternative telomere maintenance. Nature 539, 54–58 (2016).

6. O’Sullivan, R. J. & Greenberg, R. A. Mechanisms of Alternative Lengthening of Telomeres. Cold Spring Harb Perspect Biol 17, a041690 (2025).

7. Bhargava, R., Lynskey, M. L. & O’Sullivan, R. J. New twists to the ALTernative endings at telomeres. DNA Repair 115, 103342 (2022).

8. Lundblad, V. & Blackburn, E. H. An alternative pathway for yeast telomere maintenance rescues est1− senescence. Cell 73, 347–360 (1993).

9. Lafferty-Whyte, K. et al. A gene expression signature classifying telomerase and ALT immortalization reveals an hTERT regulatory network and suggests a mesenchymal stem cell origin for ALT. Oncogene 28, 3765–3774 (2009).

10. Arora, R. et al. RNaseH1 regulates TERRA-telomeric DNA hybrids and telomere maintenance in ALT tumour cells. Nature communications 5, 5220 (2014).

11. O’Sullivan, R. J. et al. Rapid induction of alternative lengthening of telomeres by depletion of the histone chaperone ASF1. Nat Struct Mol Biol 21, 167–174 (2014).

12. Flynn, R. L. et al. Alternative lengthening of telomeres renders cancer cells hypersensitive to ATR inhibitors. Science 347, 273–277 (2015).

13. Cesare, A. J. et al. Spontaneous occurrence of telomeric DNA damage response in the absence of chromosome fusions. Nat Struct Mol Biol 16, 1244–1251 (2009).

14. Thosar, S. A. et al. Oxidative guanine base damage plays a dual role in regulating productive ALT-associated homology-directed repair. Cell Reports 43, 113656 (2024).

15. Lu, R. & Pickett, H. A. Telomeric replication stress: the beginning and the end for alternative lengthening of telomeres cancers. Open Biol. 12, 220011 (2022).

16. Zhao, H. & Piwnica-Worms, H. ATR-Mediated Checkpoint Pathways Regulate Phosphorylation and Activation of Human Chk1. Molecular and Cellular Biology 21, 4129–4139 (2001).

17. Liu, S. et al. Distinct roles for DNA-PK, ATM and ATR in RPA phosphorylation and checkpoint activation in response to replication stress. Nucleic Acids Research 40, 10780–10794 (2012).

18. Silva, B. et al. FANCM limits ALT activity by restricting telomeric replication stress induced by deregulated BLM and R-loops. Nat Commun 10, 2253 (2019).

19. Pan, X. et al. FANCM suppresses DNA replication stress at ALT telomeres by disrupting TERRA R-loops. Sci Rep 9, 19110 (2019).

20. Jiang, H. et al. BLM helicase unwinds lagging strand substrates to assemble the ALT telomere damage response. Molecular Cell 84, 1684–1698.e9 (2024).

21. Wondisford, A. R. et al. Deregulated DNA ADP-ribosylation impairs telomere replication. Nat Struct Mol Biol 31, 791–800 (2024).

22. Pan, X. et al. FANCM, BRCA1, and BLM cooperatively resolve the replication stress at the ALT telomeres. Proc. Natl. Acad. Sci. U.S.A. 114, (2017).

23. Panier, S. et al. SLX4IP antagonizes promiscuous BLM activity during ALT maintenance. Molecular cell 76, 27–43 (2019).

24. Lu, R. et al. The FANCM-BLM-TOP3A-RMI complex suppresses alternative lengthening of telomeres (ALT). Nat Commun 10, 2252 (2019).

25. Sobinoff, A. P. et al. BLM and SLX4 play opposing roles in recombination- dependent replication at human telomeres. The EMBO Journal 36, 2907–2919 (2017).

26. Barefield, C. & Karlseder, J. The BLM helicase contributes to telomere maintenance through processing of late-replicating intermediate structures. Nucleic Acids Research 40, 7358–7367 (2012).

27. Stavropoulos, D. J. The Bloom syndrome helicase BLM interacts with TRF2 in ALT cells and promotes telomeric DNA synthesis. Human Molecular Genetics 11, 3135–3144 (2002).

28. Loe, T. K. et al. Telomere length heterogeneity in ALT cells is maintained by PML-dependent localization of the BTR complex to telomeres. Genes Dev. 34, 650–662 (2020).

29. Min, J., Wright, W. E. & Shay, J. W. Clustered telomeres in phase-separated nuclear condensates engage mitotic DNA synthesis through BLM and RAD52. Genes Dev. 33, 814–827 (2019).

30. Zhang, J.-M., Genois, M.-M., Ouyang, J., Lan, L. & Zou, L. Alternative lengthening of telomeres is a self-perpetuating process in ALT-associated PML bodies. Molecular Cell 81, 1027–1042.e4 (2021).

31. Karow, J. K., Constantinou, A., Li, J.-L., West, S. C. & Hickson, I. D. The Bloom’s syndrome gene product promotes branch migration of Holliday junctions. Proc. Natl. Acad. Sci. U.S.A. 97, 6504–6508 (2000).

32. Selak, N. et al. The Bloom’s syndrome helicase (BLM) interacts physically and functionally with p12, the smallest subunit of human DNA polymerase δ. Nucleic Acids Research 36, 5166–5179 (2008).

33. Zimmermann, M., Kibe, T., Kabir, S. & De Lange, T. TRF1 negotiates TTAGGG repeat-associated replication problems by recruiting the BLM helicase and the TPP1/POT1 repressor of ATR signaling. Genes Dev. 28, 2477–2491 (2014).

34. Sun, H., Karow, J. K., Hickson, I. D. & Maizels, N. The Bloom’s Syndrome Helicase Unwinds G4 DNA. Journal of Biological Chemistry 273, 27587–27592 (1998).

35. Bussen, W., Raynard, S., Busygina, V., Singh, A. K. & Sung, P. Holliday Junction Processing Activity of the BLM-Topo IIIα-BLAP75 Complex. Journal of Biological Chemistry 282, 31484–31492 (2007).

36. Wu, L. & Hickson, I. D. The Bloom’s syndrome helicase suppresses crossing over during homologous recombination. Nature 426, 870–874 (2003).

37. Raynard, S., Bussen, W. & Sung, P. A Double Holliday Junction Dissolvasome Comprising BLM, Topoisomerase IIIα, and BLAP75. Journal of Biological Chemistry 281, 13861–13864 (2006).

38. Manthei, K. A. & Keck, J. L. The BLM dissolvasome in DNA replication and repair. Cell. Mol. Life Sci. 70, 4067–4084 (2013).

39. Sobinoff, A. P. & Pickett, H. A. Mechanisms that drive telomere maintenance and recombination in human cancers. Current Opinion in Genetics & Development 60, 25–30 (2020).

40. Davies, S. L., North, P. S. & Hickson, I. D. Role for BLM in replication-fork restart and suppression of origin firing after replicative stress. Nat Struct Mol Biol 14, 677–679 (2007).

41. Root, H. et al. FANCD2 limits BLM-dependent telomere instability in the alternative lengthening of telomeres pathway. Hum. Mol. Genet. 25, 3255–3268 (2016).

42. Chang, S. et al. Multiple functions of the ALT favorite helicase, BLM. Cell Biosci 15, 31 (2025).

43. Barroso-González, J. et al. Anti-recombination function of MutSα restricts telomere extension by ALT-associated homology-directed repair. Cell Reports 37, 110088 (2021).

44. Sarkar, J. et al. SLX4 contributes to telomere preservation and regulated processing of telomeric joint molecule intermediates. Nucleic Acids Res 43, 5912–5923 (2015).

45. Chen, Y. & Yuan, J. The post translational modification of key regulators of ATR signaling in DNA replication. GENOME INSTAB. DIS. 2, 92–101 (2021).

46. Ling, C. et al. Bloom syndrome complex promotes FANCM recruitment to stalled replication forks and facilitates both repair and traverse of DNA interstrand crosslinks. Cell Discov 2, 16047 (2016).

47. Feng, S. et al. Profound synthetic lethality between SMARCAL1 and FANCM. Molecular Cell 84, 4522–4537.e7 (2024).

48. Cox, K. E., Maréchal, A. & Flynn, R. L. SMARCAL1 Resolves Replication Stress at ALT Telomeres. Cell Reports 14, 1032–1040 (2016).

49. Fielden, J. et al. Comprehensive Interrogation of Synthetic Relationships in the Human DNA Damage Response. Preprint at 10.1101/2023.08.18.553865 (2023).

50. Sirbu, B. M. et al. Identification of Proteins at Active, Stalled, and Collapsed Replication Forks Using Isolation of Proteins on Nascent DNA (iPOND) Coupled with Mass Spectrometry. Journal of Biological Chemistry 288, 31458–31467 (2013).

51. Gaggioli, V. et al. Dynamic de novo heterochromatin assembly and disassembly at replication forks ensures fork stability. Nat Cell Biol 25, 1017–1032 (2023).

52. Zwinderman, M. R. H. et al. Deposition Bias of Chromatin Proteins Inverts under DNA Replication Stress Conditions. ACS Chem. Biol. 16, 2193–2201 (2021).

53. Ozgencil, M., Dullovi, A., Christiane Higos, R. C., Hořejší, Z. & Bellelli, R. The linker histone H1–BRCA1 axis is a crucial mediator of replication fork stability. Life Sci. Alliance 6, e202301933 (2023).

54. Hanzlikova, H. et al. The Importance of Poly(ADP-Ribose) Polymerase as a Sensor of Unligated Okazaki Fragments during DNA Replication. Molecular Cell 71, 319–331.e3 (2018).

55. Vaitsiankova, A. et al. PARP inhibition impedes the maturation of nascent DNA strands during DNA replication. Nat Struct Mol Biol 29, 329–338 (2022).

56. Min, J., Wright, W. E. & Shay, J. W. Alternative lengthening of telomeres can be maintained by preferential elongation of lagging strands. Nucleic Acids Res gkw1295 (2017) doi:10.1093/nar/gkw1295.

57. Hoang, S. M. et al. Regulation of ALT-associated homology-directed repair by polyADP-ribosylation. Nat Struct Mol Biol 27, 1152–1164 (2020).

58. Ho, J. J. et al. The BLM-TOP3A-RMI1-RMI2 proximity map reveals that RAD54L2 suppresses sister chromatid exchanges. EMBO Rep 26, 1290–1314 (2025).

59. Saxena, S. & Zou, L. Hallmarks of DNA replication stress. Molecular Cell 82, 2298–2314 (2022).

60. Pladevall-Morera, D. et al. Proteomic characterization of chromosomal common fragile site (CFS)-associated proteins uncovers ATRX as a regulator of CFS stability. Nucleic Acids Research 47, 8004–8018 (2019).

61. O’Rourke, J. J., Bythell-Douglas, R., Dunn, E. A. & Deans, A. J. ALT control, delete: FANCM as an anti-cancer target in Alternative Lengthening of Telomeres. Nucleus 10, 221–230 (2019).

62. Kaya-Okur, H. S. et al. CUT&Tag for efficient epigenomic profiling of small samples and single cells. Nat Commun 10, 1930 (2019).

