## Supplementary Figures for "SLX4IP acts in parallel to FANCM to limit BLM-dependent replication stress at ALT telomeres"

### Supplementary Figure 1

Spindler et al.

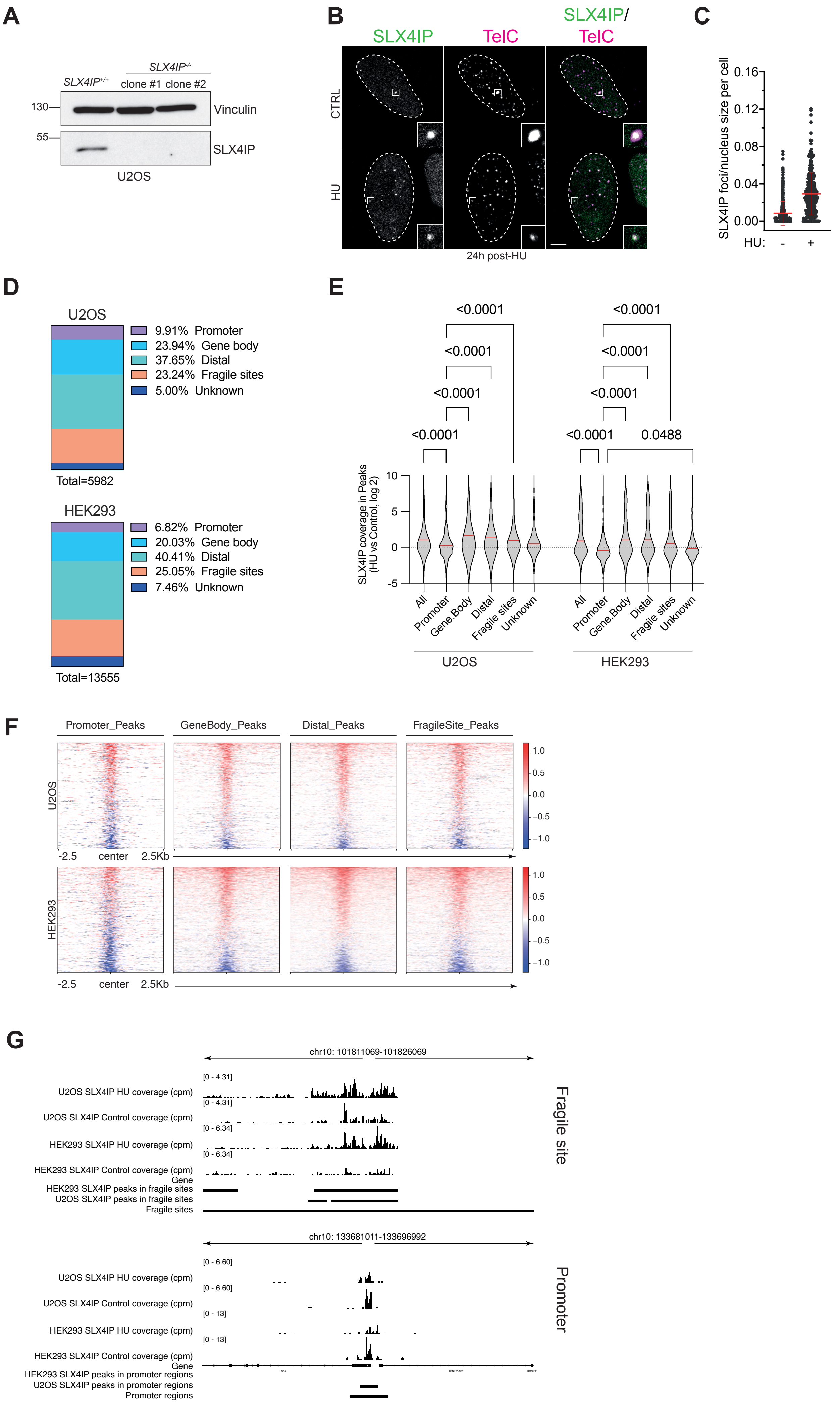

### Supplementary Figure 2

Spindler et al.

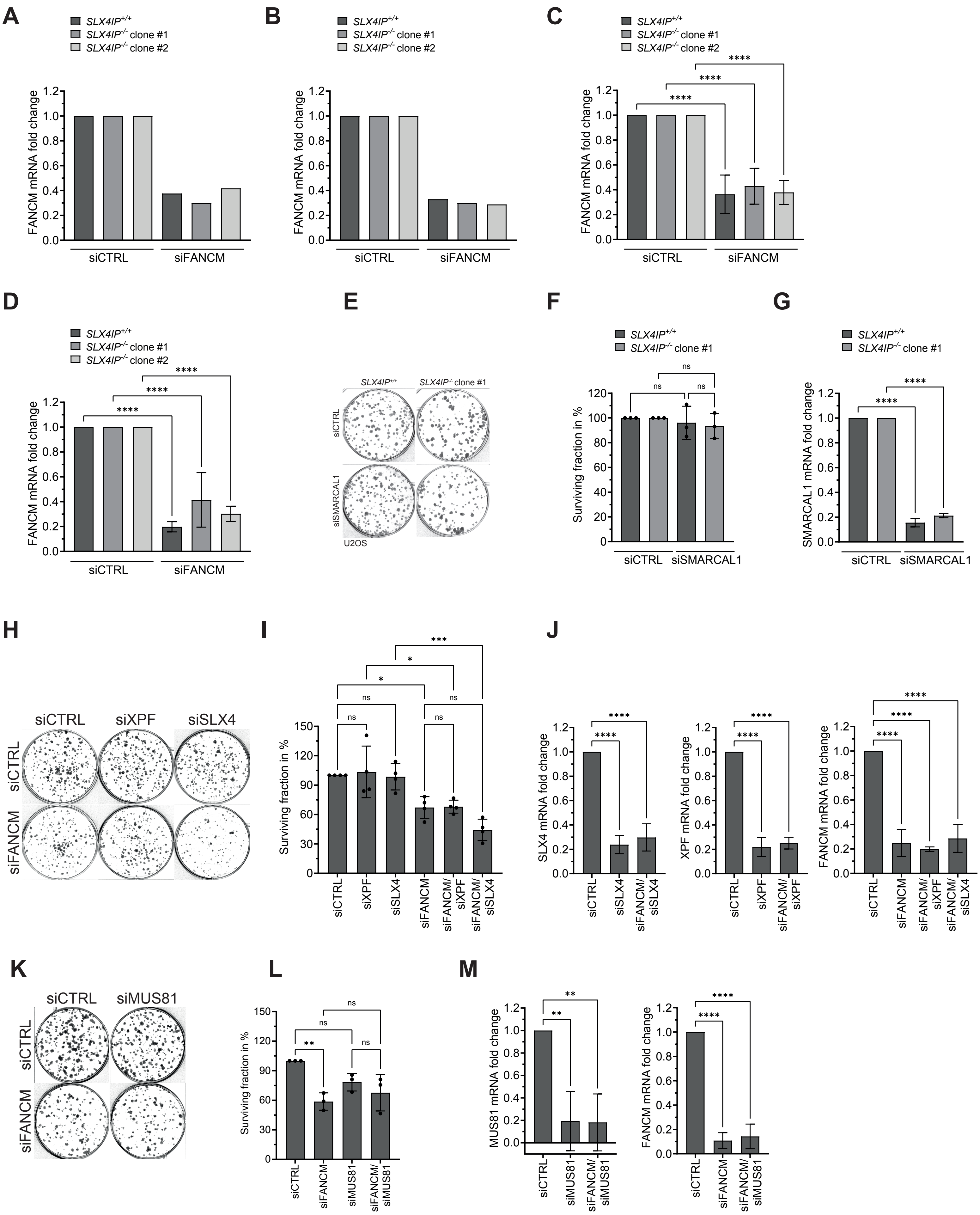

### Supplementary Figure 3

Spindler et al.

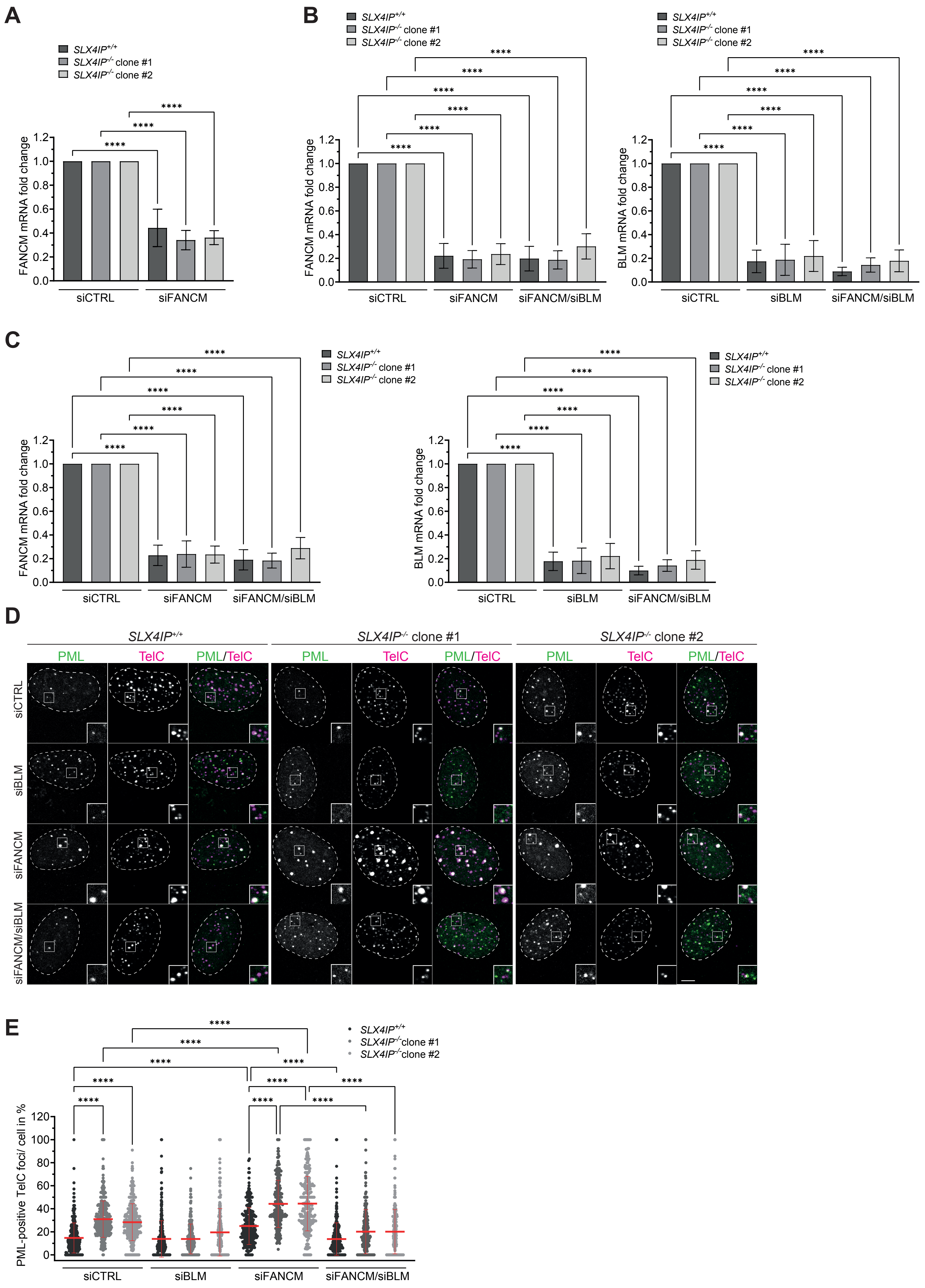

Supplementary Figure 4

Spindler et al.

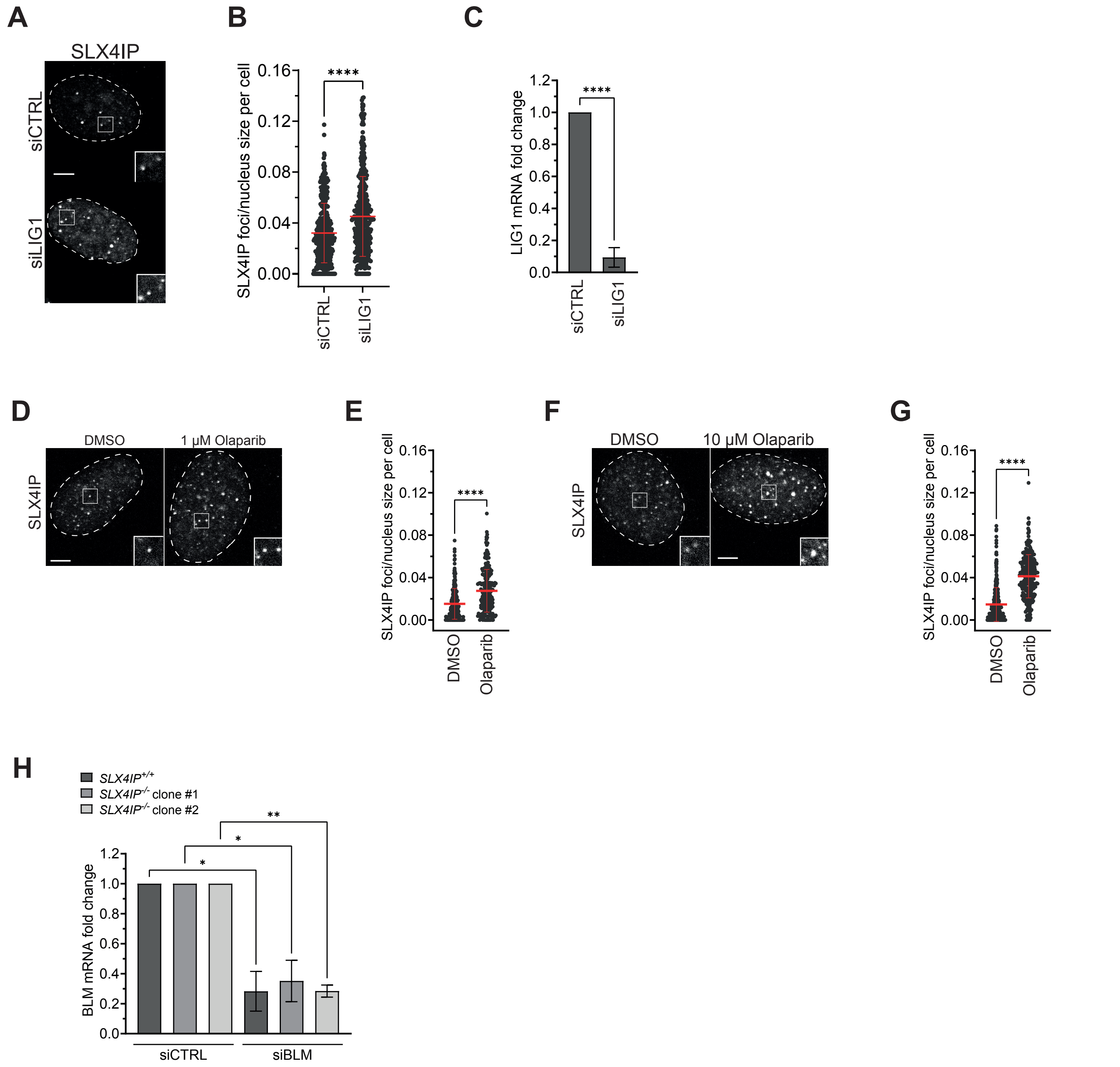
